# Single Nucleus Sequencing of Human Colon Visceral Smooth Muscle Cells, PDGFRα Cells, and Interstitial Cells of Cajal

**DOI:** 10.1101/2022.04.14.488224

**Authors:** Sabine Schneider, Sohaib K. Hashmi, A. Josephine Thrasher, Deepika R. Kothakapa, Christina M. Wright, Robert O. Heuckeroth

## Abstract

**Background and Aims:** Smooth muscle cells (SMCs), Interstitial cells of Cajal (ICCs), and PDGFRα+ cells (PαCs) form a functional syncytium in the bowel known as the ‘SIP syncytium’. The SIP syncytium works in concert with the enteric nervous system (ENS) to coordinate bowel motility. However, our understanding of individual cell types that form this syncytium and how they interact with each other remains limited, with no prior single cell RNAseq analyses focused on human SIP syncytium cells.

**Methods:** We analyzed single-nucleus RNA sequencing data from 10,749 human colon SIP syncytium cells (5572 SMC, 372 ICC, and 4805 PαC nuclei) derived from 15 individuals.

**Results:** Consistent with critical contractile and pacemaker functions and with known ENS interactions, SIP syncytium cell types express many ion channels including mechanosensitive channels in ICCs and PαCs. PαCs also prominently express ECM-associated genes and the inhibitory neurotransmitter receptor for vasoactive intestinal peptide (*VIPR2*), a novel finding. We identified two PαC clusters that differ in expression of many ion channels and transcriptional regulators. Interestingly, SIP syncytium cells co-express 6 transcription factors (*FOS*, *MEIS1*, *MEIS2*, *PBX1*, *SCMH1*, and *ZBTB16)* that may be part of a combinatorial signature that specifies these cells. Bowel region-specific differences in SIP syncytium gene expression may correlate with regional differences in function, with right (ascending) colon SMCs and PαCs expressing more transcriptional regulators and ion channels than SMCs and PαCs in left (sigmoid) colon.

**Conclusion:** These studies provide new insights into SIP syncytium biology that may be valuable for understanding bowel motility disorders and lead to future investigation of highlighted genes and pathways.

**Synopsis:** In this first single nucleus RNASeq analysis of human SIP syncytium, we identify novel features of SIP syncytium cells, including two types of PDGFRα+ cells, a SIP-specific combinatorial transcription factor signature, and colon region differences in gene expression.

## Introduction

Human bowel digests food, absorbs nutrients, eliminates waste, and protects against the entry of luminal pathogens. These processes require coordinated bowel contraction and relaxation that is mediated by cells of the “SIP syncytium” in conjunction with the enteric, sympathetic, and parasympathetic nervous system, muscularis macrophages, and enteroendocrine cells. The SIP syncytium includes visceral smooth muscle cells (SMCs) that generate force, interstitial cells of Cajal (ICCs) that act as pacemakers, and PDGFRα-expressing interstitial cells (PαCs) that modulate smooth muscle contraction and relaxation^1–3^. SMCs, ICCs, and PαCs are found in close proximity within the bowel wall and interact as functional units that collectively constitute the SIP syncytium^2^.

Visceral SMCs can produce tonic contractions to resist distension or phasic contractions to propel luminal contents^4–7^. These SMC activities are coordinated by electrically coupled ICCs that undergo spontaneous rhythmic depolarization and hyperpolarization called “slow waves”. The slow waves synchronize action potentials in SMCs to facilitate efficient propagation of muscle contractions^8^. PαCs that prominently express the cell surface receptor, PDGFRA^9^, also directly regulate smooth muscle contraction^10, 11^ and mediate purinergic inhibitory signaling^12^. SIP syncytium dysfunction can cause severe bowel dysmotility^3^.

Despite critical roles for the SIP syncytium in bowel motility, our understanding of SMC, ICCs, and PαCs phenotypes and cellular interactions remains limited. One limitation is that few studies have characterized gene expression for SMCs^13, 14^, ICCs^15, 16^, and PαCs^17^, and all published datasets are from murine intestine. We therefore analyzed our recently obtained human colon single nucleus RNAseq data^18^ to focus on the 10,749 SIP syncytium cells. These RNAseq data fit well with known mammalian physiology, but also provide new insight into SIP syncytium cell biology. Our analyses distinguished two subtypes of PαCs. Both PαC subtypes express relatively high numbers of extracellular matrix (ECM) components and remodeling proteins. One PαC cluster expresses primarily structural ECM genes, while the other cluster expresses predominantly non-structural ECM, in addition to ion channels and neurotransmitter receptors. ICCs also express many ion channels, while SMCs prominently express both ion channels and contractile apparatus constituents. One novel finding is that ICCs and PαCs express mechanosensitive ion channels, suggesting they respond directly to mechanical stimuli. PaCs also express the neurotransmitter receptor *VIPR2*, which has not been previously reported. In addition, SIP syncytium cells shared relatively high levels of 6 transcriptional regulators, suggesting a possible combinatorial transcriptional network underlying the identity of the SIP syncytium. Interestingly, gene expression differs in SMCs and PαCs from right (ascending) compared to the left (sigmoid) colon. Collectively, these analyses suggest several new ideas about how SIP syncytium cells interact and should facilitate development of new strategies to modulate the SIP syncytium to improve bowel motility.

## Methods

### Human tissue collection and single nucleus isolation

Detailed methods are in our recent manuscript^18^. Briefly, de-identified colon was acquired with Institutional Review Board (IRB) approval from the Perelman School of Medicine at the University of Pennsylvania (IRB#804376) and the Children’s Hospital of Philadelphia Institutional Review Board (IRB#13-010357) using the Abramson Cancer Center Tumor Bank.

Human colon without gross abnormalities was processed 1-4 hours after resection. Submucosa was removed before muscularis was pinned flat, stained with 4-(4-(Dimethylamino)styryl)-N-Methylpyridinium Iodide (4-Di-2-Asp; Abcam, Cat# ab145266) to visualize the enteric nervous system^18^, and carefully dissected to enrich for cells near the myenteric plexus and surrounding smooth muscle. Dissected tissue was flash-frozen in cold Optimal Cutting Temperature (O.C.T.) compound (Fisher Healthcare Tissue-Plus O.C.T. Compound; Thermo Fisher Scientific, Cat#23-730-571) and cryosectioned (100 µm sections). Tissues were only processed if RNA Integrity Number (RIN) was > 7.0, as described^18^. To isolate nuclei, frozen sections were transferred into ice-cold lysis buffer (10mM Tris HCl pH 7.5, 10 mM NaCl, 3 mM MgCl_2_, 0.005% Nonidet P40 Substitute (Thermo Fisher Scientific, Cat#AM9010; Sigma, Cat#74385)), rapidly chopped using iridectomy scissors (1 minute), transferred to a Dounce homogenizer on ice, homogenized 15 strokes with the loose pestle and 40 strokes with the tight pestle, and filtered through 30 µm MACS SmartStrainer (Miltenyi Biotec, Cat#130-098-458). Nuclei were centrifuged (590xg, 8 minutes, 4°C), resuspended in staining buffer (1x phosphate-buffered saline (PBS), 1% w/v Ultrapure BSA (Life Technologies, Cat#AM2618), 0.2U/mL Protector RNase inhibitor, (Sigma, Cat#3335399001)), and 2.5 µg/mL Hoechst 33342 Trihydrochloride Trihydrate (Thermo Fisher Scientific, Cat# H3570) and filtered again through a 40 µm FlowMi strainer (VWR, Cat#H13680-0040) before flow sorting (MoFlo Astrios) Hoechst positive nuclei into 5 µL staining buffer using a 70 µm nozzle.

### Library generation, sequencing, and data processing

Libraries were prepared using Chromium Single Cell 3’ Reagent Kits v2, (Cat#120237, 10X Genomics, Pleasanton, CA) and sequenced on an Illumina HiSeq 2500 as described^18^ (GEO accession number GSE156905). BCL to FASTQ file conversion was achieved via the Cell Ranger pipeline (10x genomics, v. 3.0.0). To aggregate data from 16 human colons, normalize outputs and re-compute gene-barcode matrices, we used the Cell Ranger Aggr pipeline (10x genomics, v. 3.0.0). Aggregated data analyzed are identical to files deposited in Gene Expression Omnibus (GEO) series GSE156905: “human_aggregated_barcodes.tsv.gz”, “human_aggregated_genes.tsv.gz”, and “human_aggregated_matrix.mtx.gz”.

### Analysis of human single-nucleus RNA sequencing data

Using Seurat^19, 20^, gene-barcode matrices were imported into R, filtered to remove low-expressors or doublets (nGene = 200-5000) and mitochondrial contaminants (percent mitochondria <10%), normalized, and scaled to regress out variance due to differing percent mitochondrial RNA and number of unique RNA molecules (UMI) identified per nucleus. Nuclei were clustered using the most statistically significant principal components up to the number where either any additional principal component contributed <5% of SD and the principal components cumulatively contributed to 90% of the SD or when the variation changes by <0.1% between consecutive principal components (17 principal components)^21^. UMAP clustering of all data allowed identification of cell clusters including PαCs, ICCs, and SMCs. PαCs were identified based on expression of *PDGFRA.* Nuclei in the ICC cluster co-expressed *KIT* and *ANO1*. Cell identity of SMCs was defined by expression of *MYH11*. To evaluate if clustering was affected by sample of origin characteristics in addition to cell type-specific gene expression, individual nuclei within the UMAP layout were color-labeled based on known sample of origin characteristics and clustering patterns were visually assessed. Sample #5035 was removed because this single sample significantly altered the clustering of SIP syncytium cell types. After removing sample #5035, the remaining data were re-normalized and re-scaled to regress out UMI and percent mitochondrial RNA. Nuclei in this smaller (n=15) dataset were clustered using the 16 most statistically significant principal components^21^. Clusters PαC#1 and PαC#2 were manually combined into a single cluster. To identify differentially expressed genes, the FindMarkers function was used to compare gene expression of SMC, ICC, and PαC clusters to that of all other nuclei in the dataset. For this and each subsequent type of differential gene expression analysis, only genes expressed by more than 10% of cells in a given cluster were included and genes enriched by more than 0.25 log_e_(fold change of mean expression level) compared to cells in all other clusters were considered to be differentially expressed.

To identify genes differentially expressed between SIP syncytium cell types, nuclei not assigned SMC, PαC, and ICC clusters were removed from the data set. Prior normalization and scaling was removed from the subsetted data. Data was then re-normalized and re-scaled to regress out UMI and percent mitochondrial RNA. Nuclei were clustered using the 10 most statistically significant principal components^21^. Nuclei separated neatly into clusters composed of SMCs, ICCs, and PαCs. Cell identity was then confirmed using canonical cell type markers for smooth muscle (*MYH11*, *ACTG2*, *ACTA2*), ICC (*KIT*, *ANO1*), and PαCs (*KCNN3*and *PDGFRA*). Differentially expressed genes between SMC, ICC, or PαC clusters compared to other SIP syncytium cell types were identified using the FindAllMarkers function. To identify gene expression differences by region of origin, nuclei within each cluster (i.e. SMC, ICC, PaC) were assigned identities of “right” or “left” colon and FindMarkers() was used to compare gene expression patterns within clusters. All p-values in this study are adjusted based on Bonferroni correction. We considered any genes with adjusted p-value <0.05 as significantly differentially expressed. All lists of differentially expressed genes generated in this study can be accessed as Supplementary Data.

### Data Visualization

Data was visualized using R and analysis was performed in GraphPad Prism version 9.3.1 for Windows (GraphPad Software, San Diego). Transcription factors were classified by “superclass” based on characteristics of their DNA-binding domains using the schema outlined by Wigender *et al.*^22^ (http://tfclass.bioinf.med.uni-goettingen.de/).

Genes enriched in the cell type of interest by greater than 0.25 log_e_(fold change of mean expression level) were submitted as gene lists to Metascape Gene Annotation and Analysis Resource for Gene Ontology and Gene Interaction Network Analysis^23^.

### Access to Data

All authors had access to the study data and reviewed and approved the final manuscript.

## Results

The single nucleus RNA sequencing data set analyzed is derived from 16 individual human sigmoid (left) or ascending (right) colon samples containing cells microdissected from peri-myenteric plexus tissue (Supplementary Table S1, and Wright and Schneider *et al*.^18^). Nuclei were isolated from homogenized tissue via fluorescence activated cell sorting and sequenced using the 10x Genomics platform (10x Genomics, Pleasanton, CA). Aggregated single nucleus data was processed and analyzed using the Seurat data visualization pipeline^20, 24^. After filtering, normalizing, and clustering, we identified 14 distinct clusters corresponding to 11 distinct cell types (Figure 1A and Supplementary Table S2), including 10,749 SIP syncytium nuclei (**5572 SMC, 372 ICC, and 4805 PαC nuclei**). Only nuclei expressing >200 unique genes (Figure 1B-C) with low mitochondrial RNA contamination (<10%) (Figure 1D) were included in analyses. This resulted in means of 1138.21 unique genes and 1983.17 unique RNAs detected per nucleus (Figure 1B-C).

**Figure 1.**
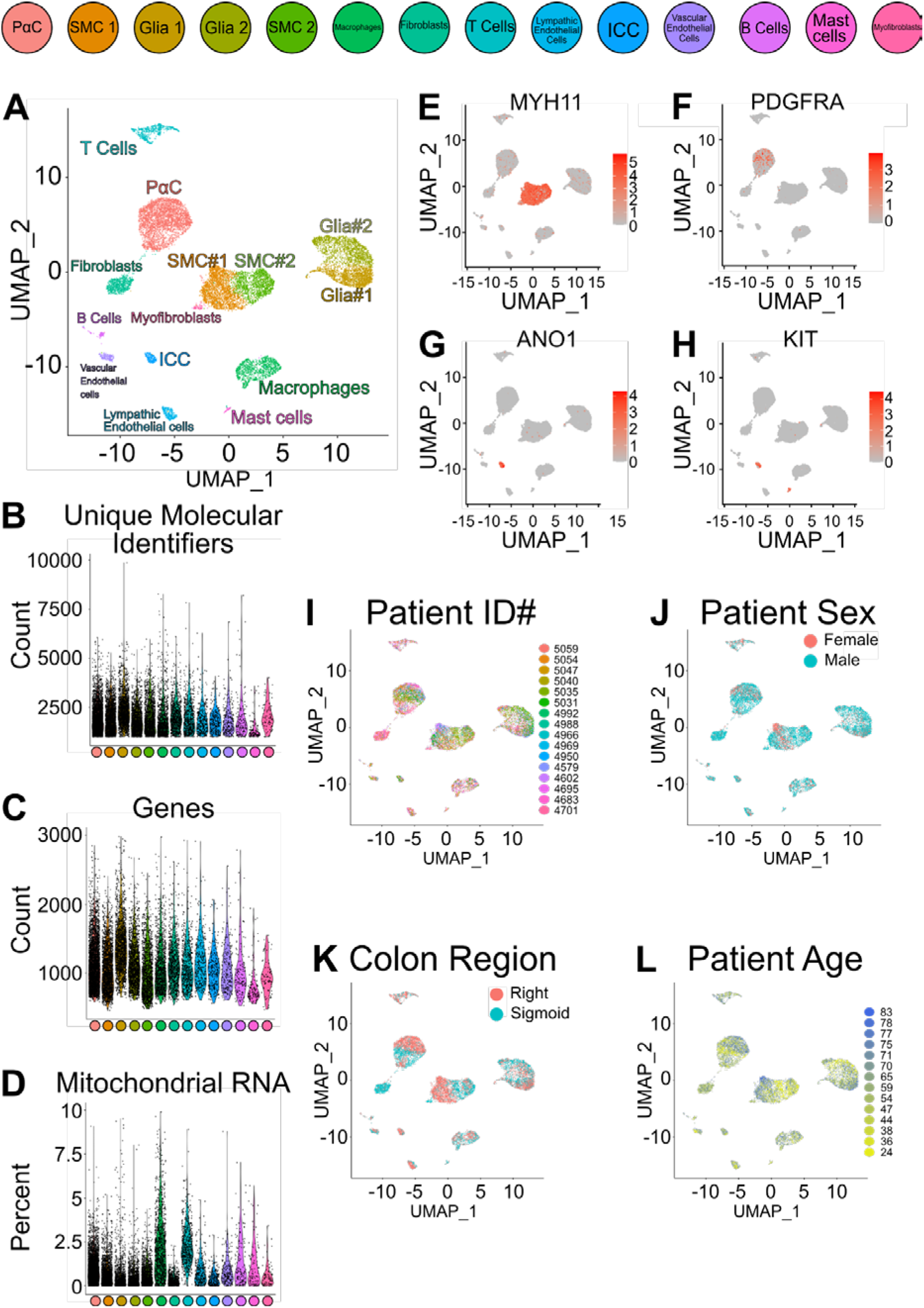
Unsupervised clustering of single nucleus RNA sequencing data from human colon muscularis reveals diverse cell types. (A) UMAP projection of all aggregated human nuclei. Each cluster of nuclei is marked by a distinct color and labeled with its putative cell type identity based on differential gene expression. SMC = smooth muscle cells, ICC = interstitial cells of Cajal, PαC = PDGFRα+ cells. (B-D) Violin plots show the distribution of the absolute number of unique molecular identifiers (unique RNA molecules) detected per cell (B), the number of unique genes identified per cell (C), and the percent mitochondrial RNA detected per cell (D). (A-D) Color legend for the clusters is provided at the top of the figure. (E-H) Featureplots visualize the differential expression of cell type-specific marker genes superimposed upon the UMAP projection shown in (A). Brighter red color indicates higher expression relative to other cell types. (E) Smooth muscle cell clusters 1 and 2 (SMC1 and SMC2) have a relatively high expression of smooth muscle myosin heavy chain 11 (*MYH11*). (F) *PDGFRA*+ cells (PαC) uniquely express the cell surface receptor *PDGFRA*. (G-H) Interstitial cells of Cajal (ICC) express relatively high levels of *ANO1* (G) and *KIT* (H), and *KIT* is also expressed by mast cells. Relative expression is given as log_e_(expression level in cell/mean expression level across all clusters). (I-L) UMAP projection from panel (A) with specific characteristics of the 16 aggregated samples highlighted. (I) Nuclei from each human biospecimen are identified by a distinct color. Sample ID numbers correspond to individuals in Supplementary Table 1. (J) Male and female nuclei are color coded. (K) Nuclei are colored by colon region of origin. (L) Nuclei are color coded by age in years.

Unsupervised clustering of all nuclei yielded two SMC clusters (SMC#1 and SMC#2) with prominent expression of smooth muscle myosin heavy chain 11 *MYH11* (Figure 1A, E), one PαC cluster, identified based on expression of *PDGFRA* (Figure 1F), and one ICC cluster based on *ANO1* and *KIT* expression (Figure 1G-H). Reassuringly, nuclei from individual samples were distributed across the 14 clusters (Figure 1I). Initial analyses suggested sex (Figure 1J) and colon region of origin (right/ascending colon versus left/sigmoid colon, Figure 1K) might determine SIP syncytium cell type clustering patterns, whereas there was no effect of age (Figure 1L). However, omitting a single sample (#5035) from a 24-year old male with volvulus significantly changed SIP syncytium nuclei clustering patterns (Figure 2A-C). Since volvulus causes bowel ischemia and omitting other samples did not change clustering patterns, we decided to continue our analyses excluding sample #5035 (final analyses using 2976 SMC nuclei, 233 ICC nuclei, and 3117 PαC nuclei). When #5035 is omitted, there is still one ICC cluster, but two distinct PαC clusters, and only a single SMC cluster (Figure 2C). Excluding #5035 also minimized sex and colon region of origin effects on unsupervised clustering patterns (Figure 2D-E).

**Figure 2.**
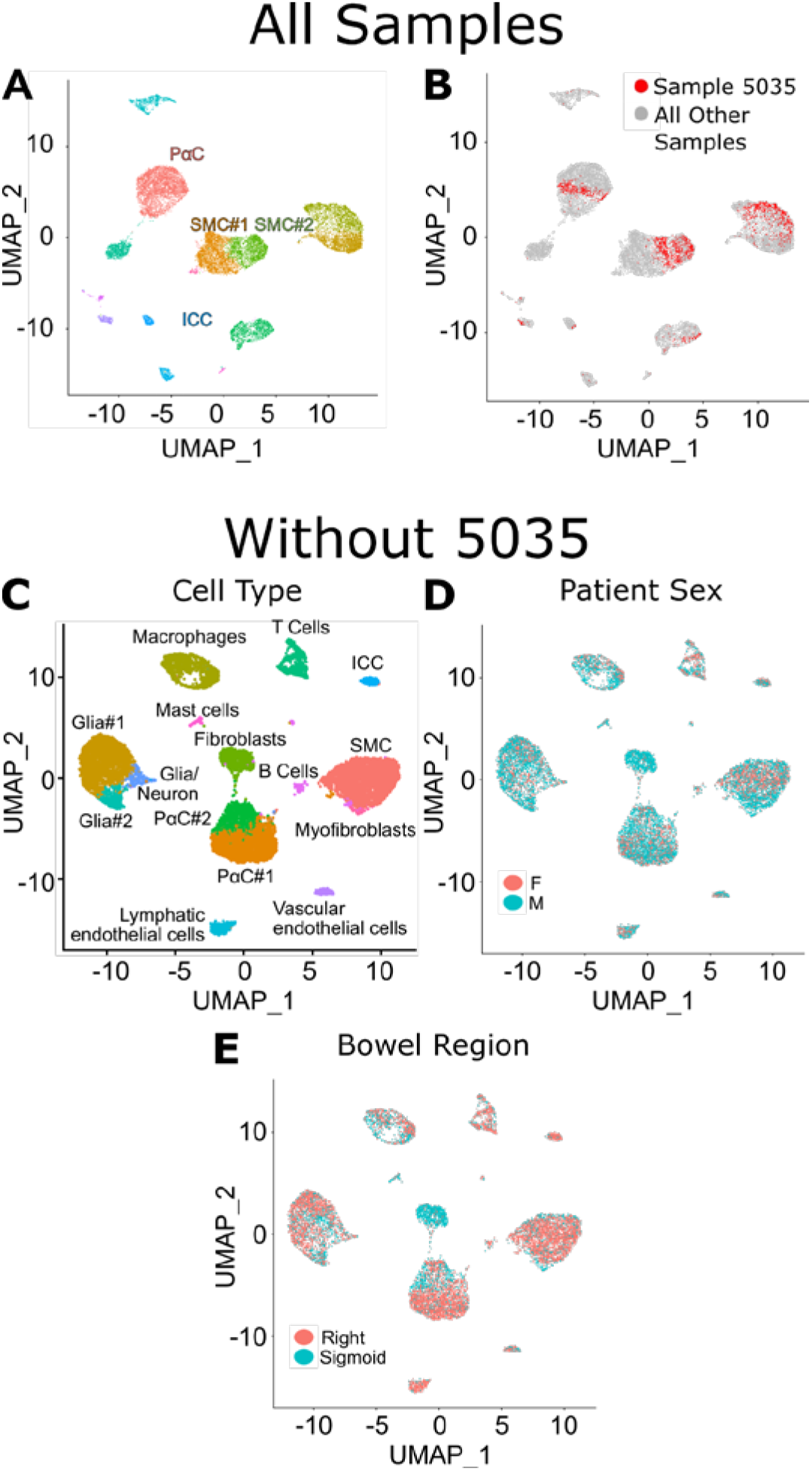
A single sample (#5035) is a major driver for the initial clustering pattern of SIP syncytium cells and was excluded from further analysis. (A) UMAP projection of the complete dataset with all aggregated human nuclei. Clusters of SIP syncytium cell types are labeled. (B) UMAP projection in panel (A) with only the nuclei derived from sample #5035 colored red. (C) UMAP projection of aggregated human nuclei derived from 15 samples, excluding sample #5035. (D) UMAP projection from panel (C) showing the distribution of nuclei derived from males versus females. (E) UMAP projection shown in (C) with nuclei colored according to colon region of origin (right/ascending colon versus left/sigmoid colon). (A, C) Smooth muscle cells (SMC, SMC#1 or SMC#2), interstitial cells of Cajal (ICC), and PαC cells (PαC, PαC#1 or PαC#2).

### Comparing gene expression profiles of SIP syncytium cell types

After omitting #5035, we re-confirmed cluster classification using *ACTG2*, *MYH11*, and *ACTA2* in the SMC cluster (Figure 3A-C), *KIT* and *ANO1* in the ICC cluster (Figure 3D-E), and *PDGFRA*, *KCNN3*, and *P2RY1* (Figure 3F-H)^2, 17^ in the two PαC clusters. To facilitate comparisons, we manually combined PαC#1 and PαC#2 into a single cluster (PαC). We then identified mRNA relatively more abundant in SIP syncytium cells compared to other cells in our data set and from this enriched gene list we identified mRNA differentially expressed between SMCs, ICCs, and PαCs. We focused on ECM components, ECM remodeling proteins, ion channels, axon guidance proteins, and synaptic proteins, and transcriptional regulators (Figure 3I-K). These categories were selected because of known functions of SIP syncytium cells. SMCs and ICCs express more mRNA for synapse associated proteins, axon guidance proteins, ion channels and ion channel-associated proteins than PαCs. (Figure 3I-K). In contrast, PαCs express more ECM components and ECM remodeling protein mRNA than SMCs or ICCs (Figure 3I-K).

**Figure 3.**
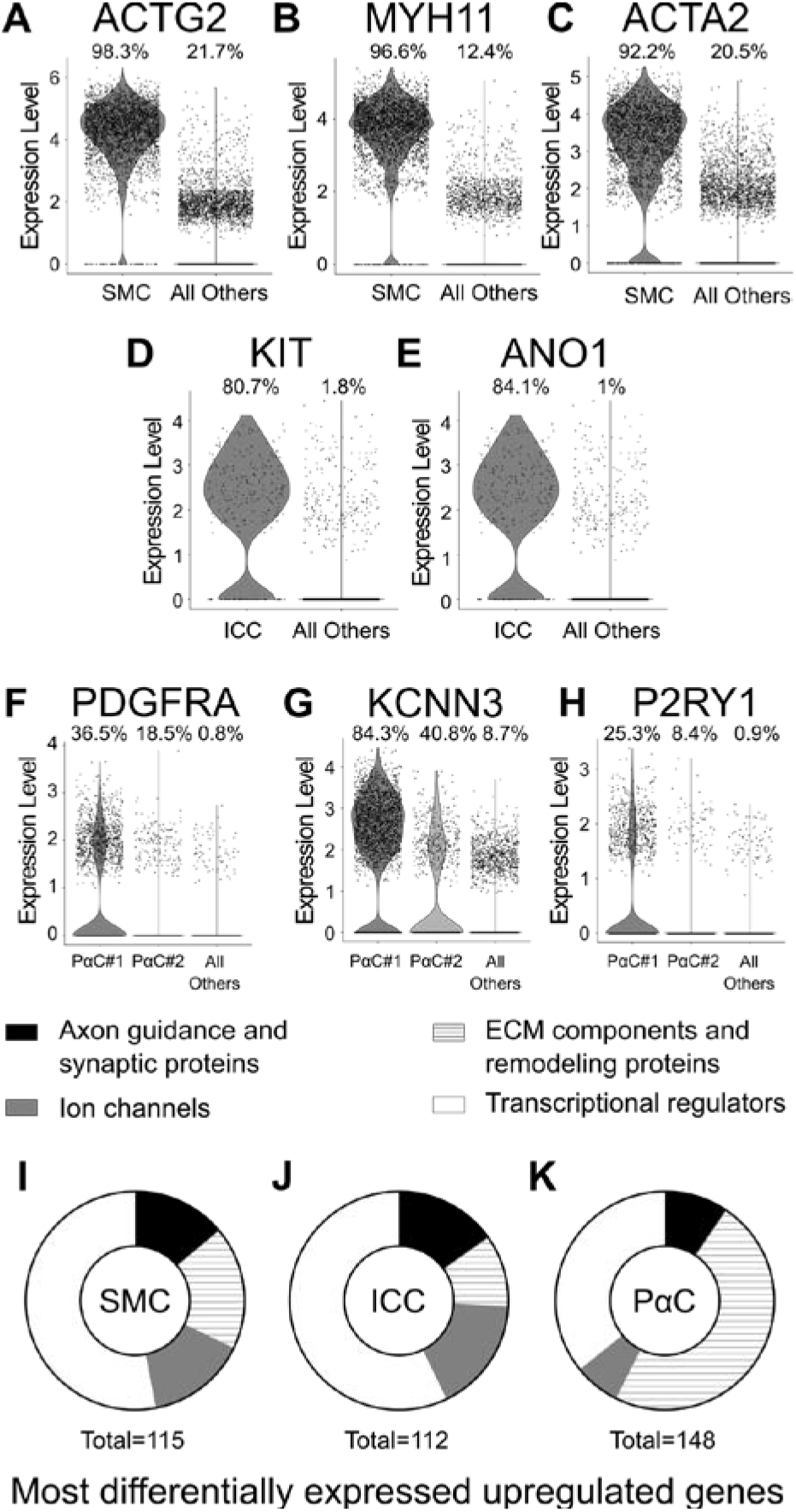
The dataset contains clusters of nuclei that correspond to SMCs, ICCs, and PαCs with distinct gene expression profiles. (A-C) The putative SMC cluster expresses relatively high levels of canonical visceral smooth muscle markers *ACTG2* (A)*, MYH11* (B), and *ACTA2* (C). (D, E) The ICC cluster expresses relatively high levels of the cell type markers *KIT* (D) and *ANO1* (E). (F-H) The two PαC clusters express the canonical PαC markers *PDGFRA* (F), *KCNN3* (G), and *P2YR1* (H). Transcripts encoding these PαC markers were detected in a greater percentage of cells in cluster PαC#1 than in PαC#2. (I-K) Using only unique genes differentially enriched in SMC, ICC, and PαC compared to all other cells in our dataset, pie charts illustrate proportions of identified genes encoding ECM components and remodeling proteins, ion channels, axon guidance and synaptic proteins, and transcriptional regulators enriched in SMC (I), ICC (J), and PαC (K). Numbers below pie charts indicate the number of genes represented in the pie chart. (A-H) Violin Plots show gene expression level for individual nuclei as a single data point laid over the proportional distribution of gene expression levels for all nuclei in the cluster. Expression level is defined as log_e_(expression level in nucleus/mean expression level for nuclei across all clusters). Proportion of nuclei in each cluster expressing the gene of interest is shown above each violin plot.

Network analysis for GO term protein-protein interaction enrichment using Metascape^23^ (Figure 4) confirmed that PαCs expressed a greater number of ECM components compared to all other cell types (Figure 4A). ICCs share many abundant, differentially expressed genes with SMCs and PαCs (Figure 4B). However, SMCs and PαCs share very few of the abundant, differentially expressed genes (Figure 4B). The most significantly enriched protein-protein interaction networks by MCODE analysis show SMCs express gene networks involved in focal adhesions, axon and neuron projection guidance, TGFβ signaling, and non-structural ECM components (Figure 5A). ICCs expressed many gene networks implicated in neurotransmission (Figure 5B) while the most highly differentially expressed gene networks in PαCs are involved in ECM production, axon guidance and nitrergic signaling (Figure 5C).

**Figure 4:**
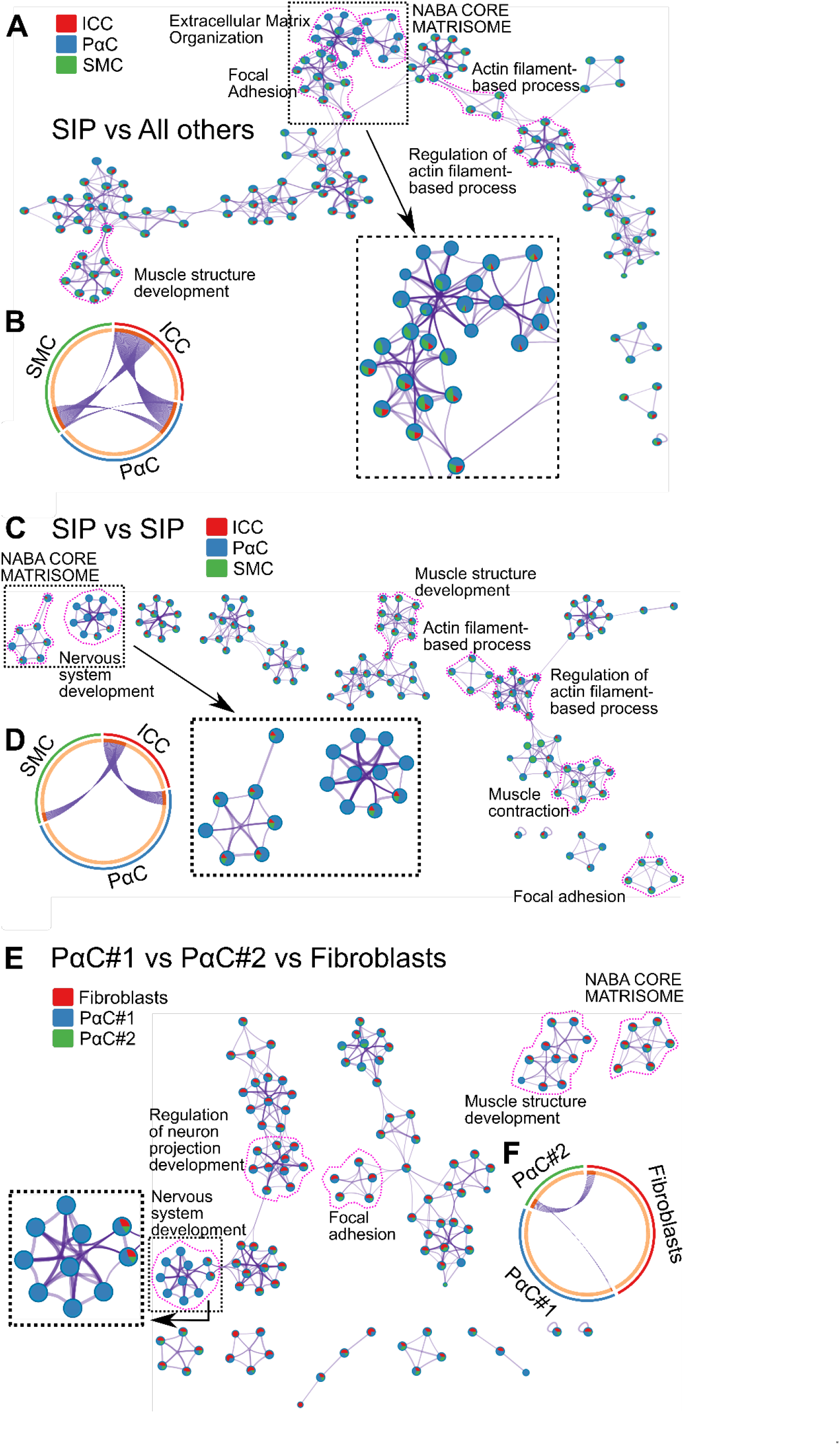
Metascape protein-protein Interaction Network Analysis. (A,C,E) The most highly enriched GO Terms from Metascape analysis are visualized as a network plot where each node is a pie chart that represents individual GO Terms and similar GO Terms are connected by edges. Relative sizes of pie chart slices correspond to the percentage of total GO Term-related genes that are differentially more abundant in each cell type of interest: (A) Based on genes enriched in the three SIP syncytium cell types (SMC, ICC, and PαC) compared to all other cells in the dataset. Green = SMC, blue = PαC, and red = ICC. (C) Based on genes enriched in each SIP syncytium cell type compared to the two other SIP syncytium cell types. Green = SMC, blue = PαC, and red = ICC. (E) Based on genes enriched in PαC#1, PαC#2, or fibroblast clusters when these cell types are compared to each other. Red = Fibroblasts, blue = PαC#1, and green = PαC#2. (B,D,F) Metascape Circos plots visualizing differentially enriched genes shared by the cell types of interest. The inner circle indicates gene lists differentially enriched in each cell type. Genes unique to a cell type are shown in light orange and genes shared by more than one cell type are shown in dark orange and are connected by purple curves. (B) Shared genes enriched in the three SIP syncytium cell types (SMC, ICC, and PαC) when gene expression is compared against all other cell types in the dataset. (D) Shared genes enriched in the three SIP syncytium cell types (SMC, ICC, and PαC) when gene expression of each SIP syncytium cell type is compared against the two other SIP syncytium cell types. (F) Shared genes enriched in each of the three fibroblast-like cell types (PαC#1, PαC#2, and Fibroblasts) when gene expression is compared against that of the other two fibroblast-like cell types. (A, C, E) Selected regions of the network are enlarged to show color coding on pie charts.

**Figure 5:**
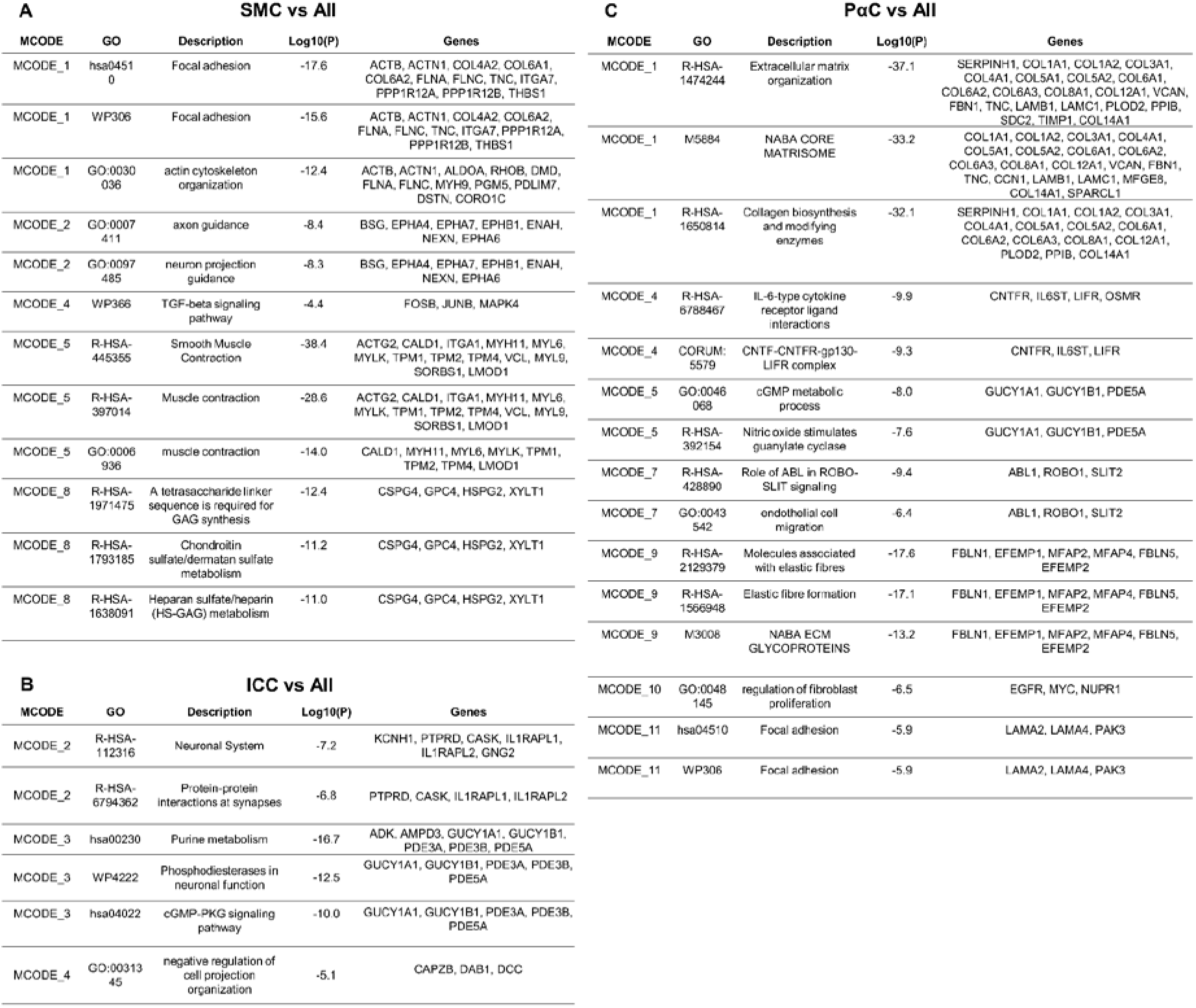
Protein-protein Interaction Enrichment Analysis - Comparing differential gene expression of SIP syncytium cell types against all other cell types in the dataset. (A-C) Protein-protein interaction networks composed of >3 differentially overexpressed genes for each SIP syncytium cell type were subjected to MCODE process enrichment analysis using Metascape. Representative GO Terms corresponding to densely connected protein-protein network components are shown for SMCs (A), ICCs (B), and PαCs (C). Differentially expressed genes were determined by comparing gene expression of each of the SIP syncytium cell types against that of all other cell type clusters in our dataset.

### SMCs express genes that facilitate interactions with the enteric nervous system

As expected, genes most abundantly and differentially expressed in the SMC cluster compared to all other cell types are canonical smooth muscle genes (*ACTG2, MYH11, TAGLN, ACTA2, MYL9, TPM2, CNN1*) involved in the SMC contractile apparatus. Several ion channels (*CACNA1C, KCNMA1, RYR2, CACNB2, KCNAB1*) and cholinergic G-protein coupled receptors (GPCRs) (*CHRM2, CHRM3)* are also much more abundant in the SMC cluster compared to all other cell types (Figure 6A). In addition, nuclei of the SMC cluster express many genes at relatively high levels that would facilitate interaction with the enteric nervous system (ENS). For example, compared to other clusters, SMCs express relatively high levels of *GPM6A* (Fold change 5.55), a glycoprotein involved in neuronal differentiation (24), *SEMA3A* (Fold change 2.79), a guidance molecule important for neuronal patterning, *NEGR1* (Fold change 2.76), a neuronal cell adhesion molecule, *NLGN1* (Fold change 2.56), which promotes glutamatergic synapse formation, and *NAV2* (Fold change 2.42), a neuron navigator. *GPM6A* expression has not previously been reported in SMCs.

**Figure 6.**
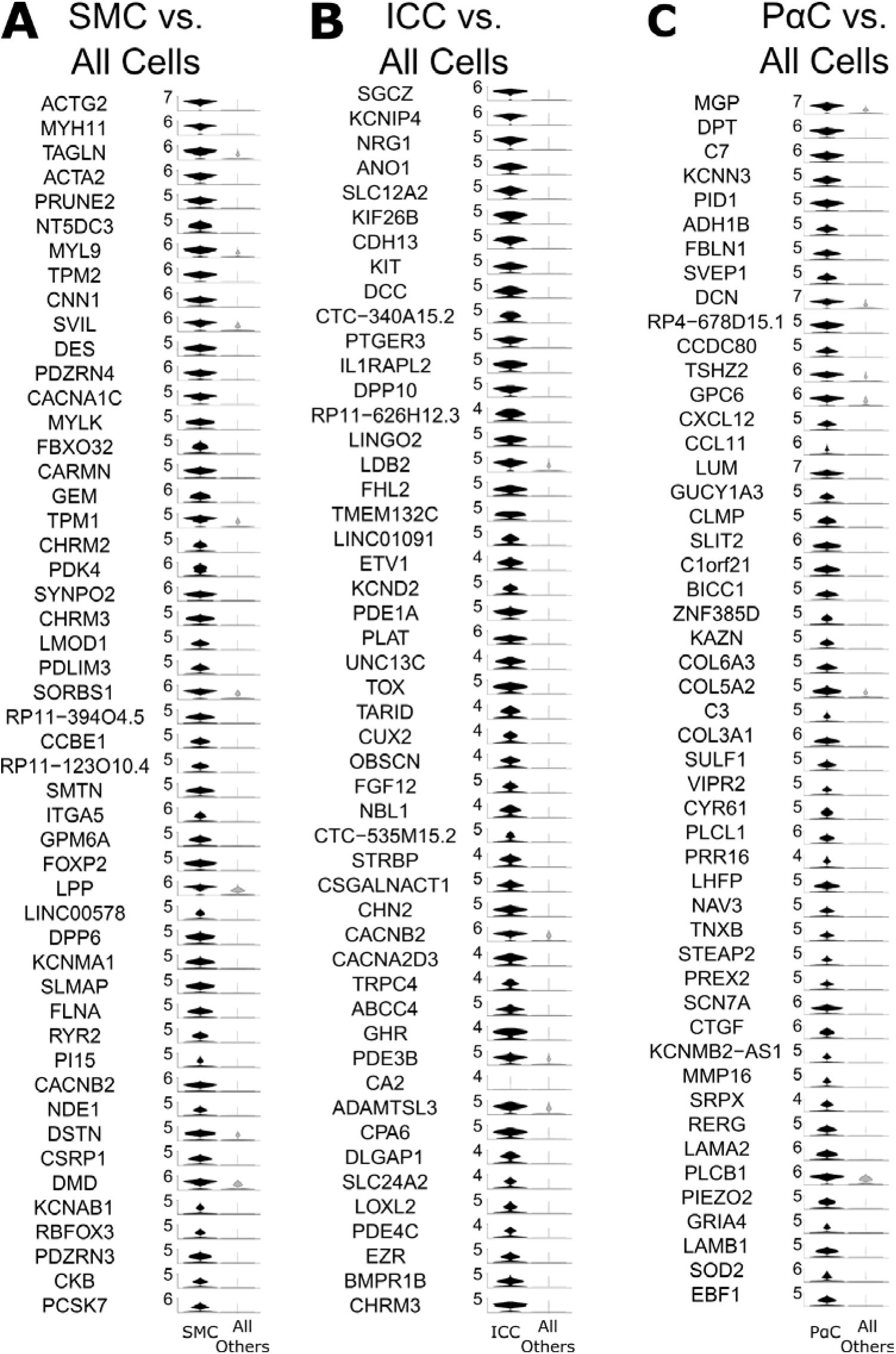
Most highly expressed genes differ for SMCs, ICCs, and PαCs. Violin plots of the top 50 most differentially expressed genes with transcripts more abundant in (A) SMCs, (B) ICCs, and C) PαCs (combined cluster including nuclei from both PαC#1 and PαC#2 clusters) compared to all other cells in our dataset. (A-C) Violin Plots show distribution of gene expression levels per individual nucleus. Expression level is defined as log_e_(expression level in nucleus/mean expression level for nuclei across all clusters).

### ICCs express a mechanosensitive ion channel, PIEZO2

The ICC cluster expresses relatively high levels of selected ion channels (*KCNIP4*, *ANO1*, *KCND2*, *CACNB2*, *CACNA2D3*, *TRPC4*) and the cholinergic muscarinic receptor *CHRM3* (Figure 6B). We also identified many highly differentially expressed genes that were relatively abundant in the ICC cluster but were not previously reported in ICCs. This list includes potassium voltage-gated channel interacting protein 4, *KCNIP4* (Fold change 21.4), sarcoglycan zeta, *SCGZ* (Fold change 21.6), cadherin 13, *CDH13* (Fold change 11.4), DCC Netrin 1 receptor, *DCC* (Fold change 10.4), and Piezo Type Mechanosensitive Ion Channel Component 2, *PIEZO2* (Fold change 3.36).

### PαCs express high levels of ECM components and mechanosensitive ion channels

The combined PαC cluster prominently expresses ion channels (*KCNN3*, *SCN7A*, *PIEZO2*, and *GRIA4*). While *PDGFRA* was not within the top 50 most differentially expressed genes in the PαC cluster, another well-known PαC marker, *KCNN3*, is the third-most differentially expressed gene in the PαCs compared to all other cells. Interestingly, the PαC cluster highly differentially expressed the neurotransmitter receptor *VIPR2* (Fold change 4.32), which has not previously been reported in PαCs, and relatively high levels of *GUCY1A3* (Fold change 5.23), which is activated by nitric oxide (Figure 6C), suggesting nitric oxide directly signals in human PαCs. Interestingly, the PαC cluster also has high expression for many genes encoding ECM components or ECM-remodeling proteins, which account for 16 of the top 50 most differentially expressed genes (Figure 6C). These genes include *MGP*, *DPT*, *FBLN1*, *SVEP1*, *DCN*, *GPC6*, *LUM*, *KAZN*, *COL6A3*, *COL5A2*, *COL3A1*, *SULF1*, *TNXB*, *MMP16*, *LAMA2*, and *LAMB1*. Furthermore, compared to SMC and ICC, PαC cluster cells express high levels of structural ECM (collagens and versican), nonstructural ECM (such as fibronectin, laminin, proteoglycans)^25^, and transcripts encoding ECM remodelers (such as matrix metalloproteinases; Figure 7A-C). The relative proportions of structural and nonstructural ECM components and ECM remodeling proteins are shown in Figure 7D.

**Figure 7:**
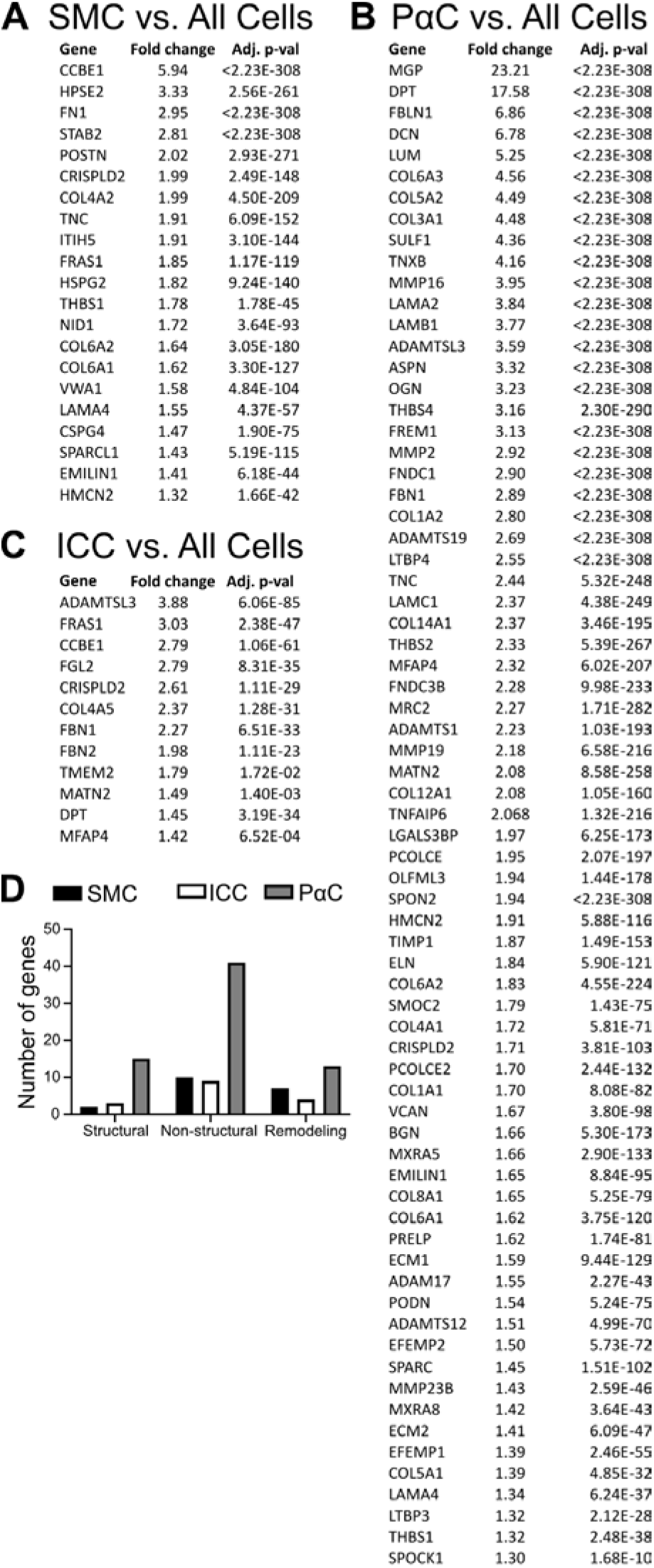
PαCs produce more ECM components than SMCs or ICCs. The most differentially expressed ECM components and ECM remodeling genes that are more abundant in (A) SMCs, (B) PαCs, and (C) ICCs compared to all other clusters. (D) PαCs express relatively high levels of a greater number of total ECM components and ECM remodelers than ICCs and SMCs. PαCs also express relatively high levels and a greater proportion of structural ECM proteins than either ICCs or SMCs. (A-C) Adj. p-val = Bonferroni adjusted p-value for each gene. Fold Change = mean number of transcripts in cluster of choice/mean number of transcripts in cells from all other clusters combined.

### SIP syncytium cells express diverse ion channels

All SIP syncytium cell types express many ion channels and ion channel-associated proteins, including subunits of potassium channels, calcium channels, sodium channels, and glutamate ionotropic receptors (Figure 8A-C). Interestingly, ICC and PαC clusters differentially express relatively high levels of the mechanosensitive channels (*PIEZO2* in ICC and PαC, and *TMEM150C* in ICC; Figure 8B and C). A variety of axon guidance and synapse-associated proteins are also enriched in SIP syncytium cells (Figure 9A-C).

**Figure 8:**
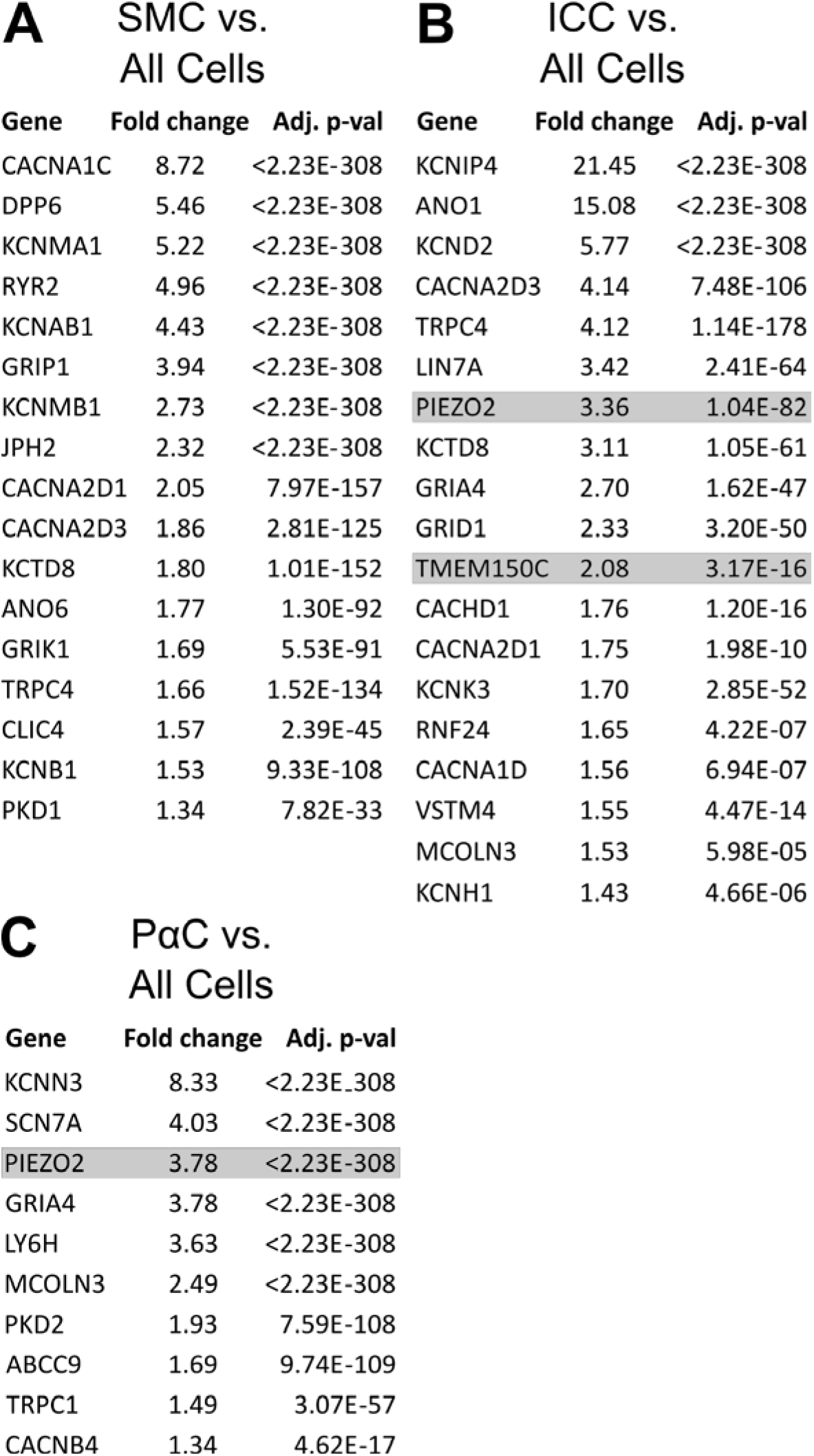
ICCs and PαCs express many ion channels, including mechanosensitive ion channels. The most differentially expressed ion channels and ion channel-associated proteins found to be more abundant in (A) SMCs, (B) ICCs, and (C) PαCs when compared to cells in all other clusters. ICCs and PαCs express mechanosensitive channels *PIEZO2* and *TMEM150C* (gray highlight). (A-C) Adj. p-val = Bonferroni adjusted p-value is shown for each gene. Fold Change = mean number of transcripts in cluster of choice/mean number of transcripts in cells from all other clusters combined.

**Figure 9.**
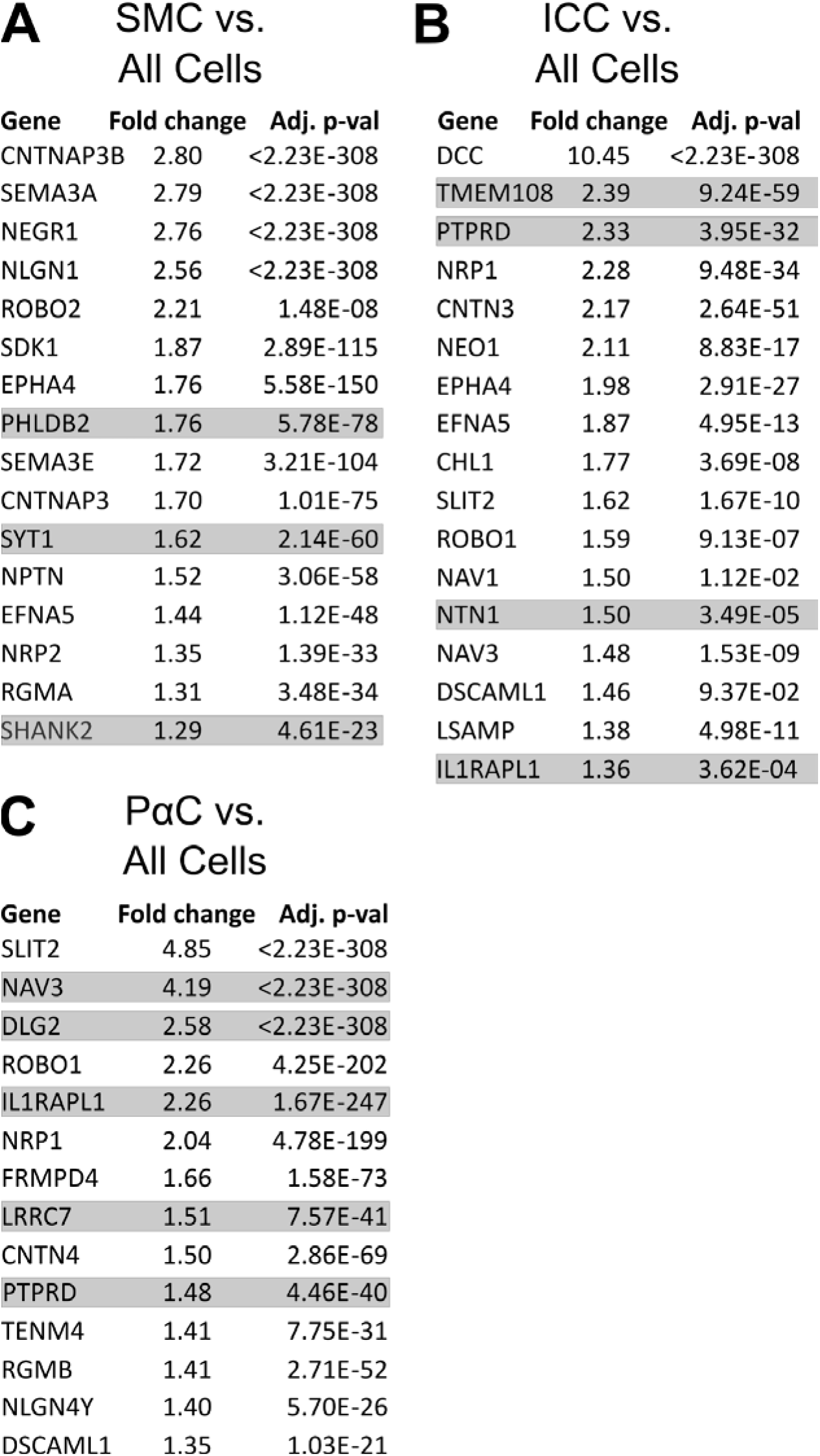
Transcripts encoding proteins involved in axon guidance and synapse function/maintenance are differentially expressed at relatively high levels in SIP syncytium cells. The most differentially expressed axon guidance and synapse-associated proteins expressed more abundantly in (A) SMCs, (B) ICCs, and (C) PαCs when compared to cells in all other clusters. (A-C) Synapse-associated proteins are highlighted in gray. Adj. p-val = Bonferroni adjusted p-value is shown for each gene. Fold Change = mean number of transcripts in cluster of choice/mean number of transcripts in nuclei from all other clusters combined.

### SIP syncytium shares a unique signature of 6 transcriptional regulators

SMCs, ICCs, and PαCs each differentially express at relatively high levels many transcriptional regulators (including epigenetic regulators, direct transcriptional activators, and repressors; Figure 10A-C). A subset of these transcriptional regulators was expressed at relatively high levels by all three SIP syncytium cell types compared to all other clusters in the dataset (Figure 11A). This shared subset includes *FOS*, *MEIS1*, *MEIS2*, *PBX1*, *SCMH1*, and *ZBTB16* (Figure 11B). A diversity of transcription factor classes are expressed by all SIP syncytium cells (Figure 11C-E).

**Figure 10.**
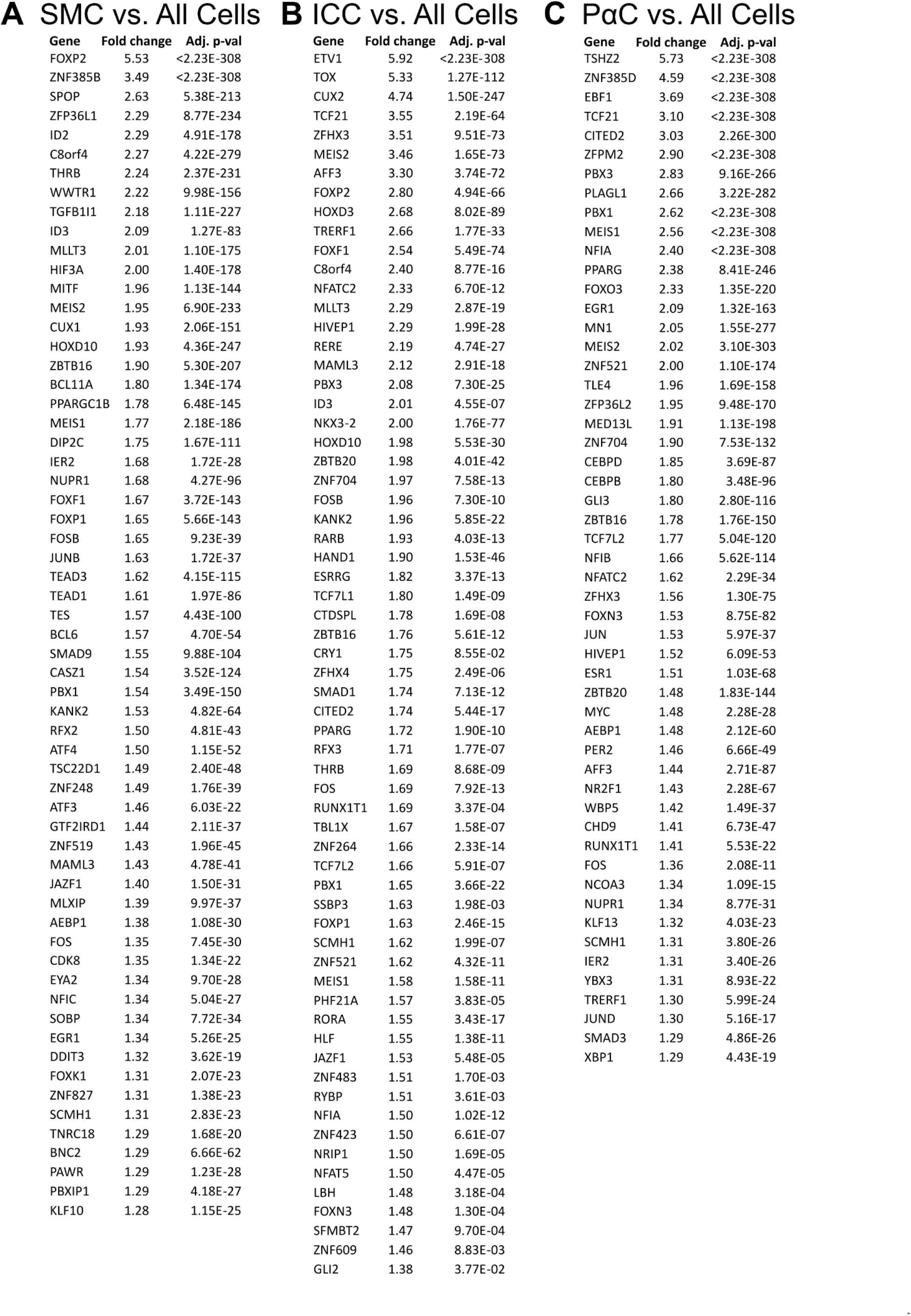
SIP syncytium cells differentially express a large number of transcriptional regulators. A) SMCs, B) ICCs, and C) PαCs differentially express similar numbers of transcriptional regulators at increased levels compared to all other cells in our dataset. (A-C) Adj. p-val = Bonferroni adjusted p-value is shown for each gene. Fold Change = mean number of transcripts in cluster of choice/mean number of transcripts in cells from all other clusters combined.

**Figure 11:**
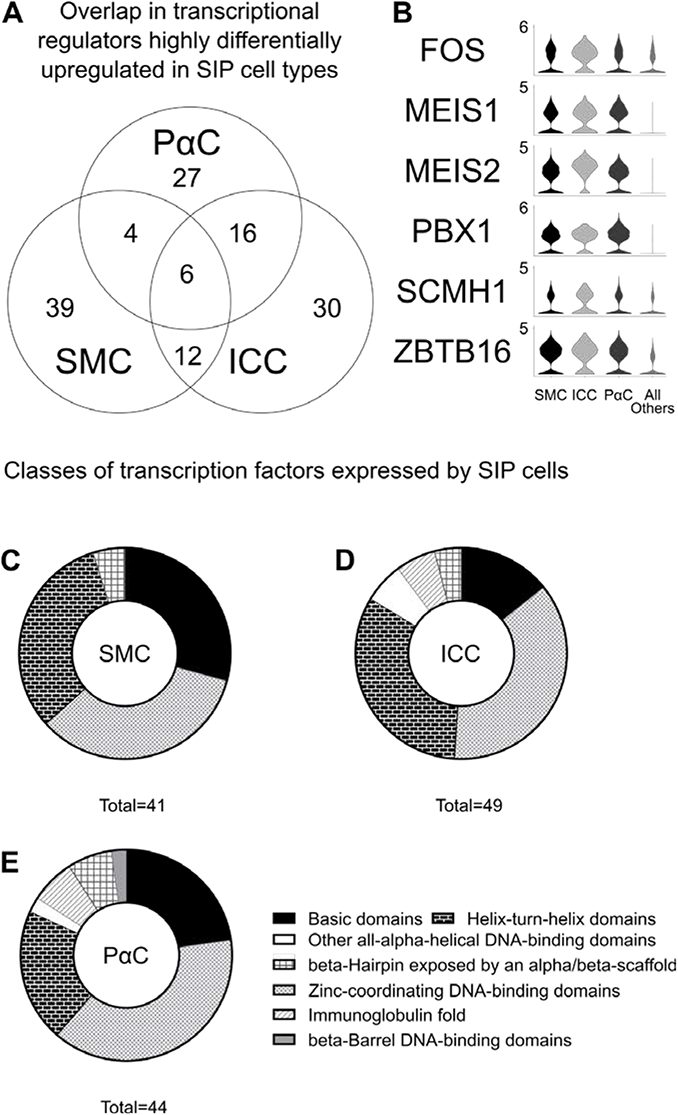
SIP syncytium cells express many transcriptional regulators and a subset are shared among SIP syncytium cell types. (A) Venn diagram shows co-expression of transcriptional regulators expressed at relatively high levels in SIP syncytium cells compared to all other cells in the dataset. (B) Violin plots show relative expression of the 6 transcriptional regulators co-expressed at relatively high levels by all SIP syncytium cell types. (C-E) Many transcriptional regulator classes are expressed by SIP syncytium cell types. Numbers below pie charts indicate the number of genes represented in the pie chart.

### Compared to other SIP syncytium clusters, PαCs abundantly express transcriptional regulators and ECM components

For the preceding analyses, gene expression in SIP syncytium cells was compared to all other cells in our dataset. This allowed us to determine how SIP syncytium cell types differ from other cell types. However, this global comparison could mask more subtle differences between SIP cell types themselves. We therefore removed non-SIP syncytium clusters from the dataset, and re-normalized and re-scaled our data. We again identified many differentially expressed genes for ion channels, ion channel-associated proteins, axon guidance and synaptic proteins, as well as ECM components and remodeling proteins (Figure 12A-C). Curiously, structural ECM mRNA were relatively more abundant in PαCs cells, whereas SMCs and ICCs expressed proportionally more ECM remodelers (Figure 12D-F).

**Figure 12:**
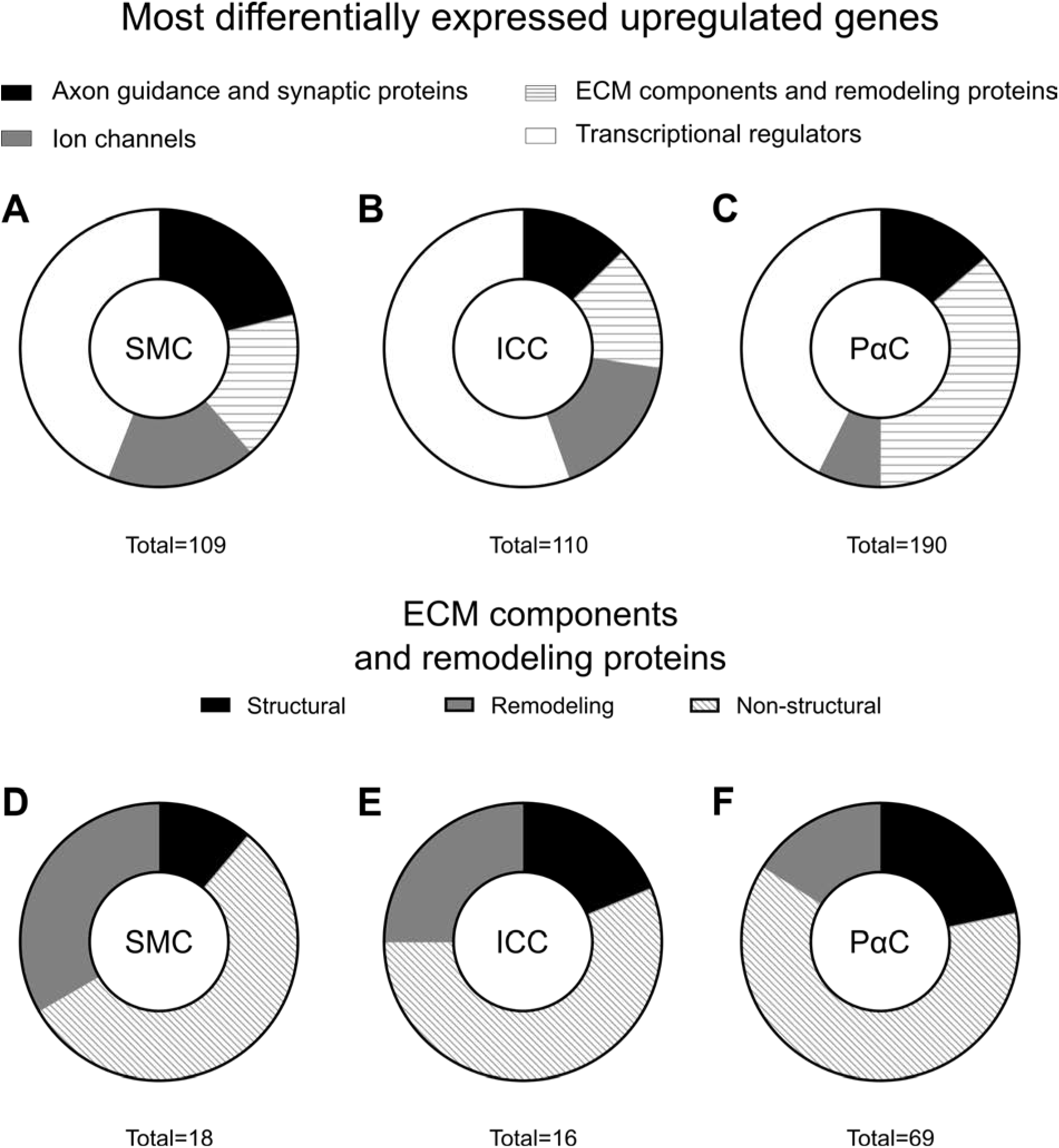
SIP syncytium cell types differentially express axon guidance and synaptic proteins, ion channels, ECM components and transcriptional regulators. (A-C) Pie charts illustrate the relative proportion of differentially expressed transcripts encoding ECM components and remodeling proteins, ion channels, axon guidance and synaptic proteins and transcriptional regulators that are more abundant in SMCs (A), ICCs (B), and PαCs (C) when compared only to other SIP syncytium cells. (D-F) PαCs express ∼4 times as many ECM associated genes as SMCs and ICCs. Pie charts illustrate the relative proportion of differentially more abundant genes encoding structural and nonstructural ECM components as well as ECM remodelers in SMCs (D), ICCs (E), and PαCs (F) when compared only to other SIP syncytium cell types. Numbers below pie charts indicate the number of genes represented in the pie chart.

Metascape network analysis comparing SIP nuclei to each other (Figure 4C) highlighted many of the same GO term networks identified when SIP syncytium cells were compared to all other cells in our dataset (Figure 4B). SMC and ICC share a subset of differentially enriched mRNA transcripts, whereas ICC and PαCs share a non-overlapping set of differentially enriched mRNA. However, SMCs and PαCs have no differentially enriched mRNA transcripts in common when we analyze only SIP syncytium cells (Figure 4D). MCODE protein-protein interaction networks show axon guidance pathways are enriched in SMCs compared to other SIP cell types (Figure 13A). Analysis of ICCs showed additional networks involved in neurotransmission beyond those identified by the comparison against all other cell types (Figure 13B). Surprisingly, there were no MCODE enrichment terms identified for the PαC when compared only to other SIP cell types.

**Figure 13:**
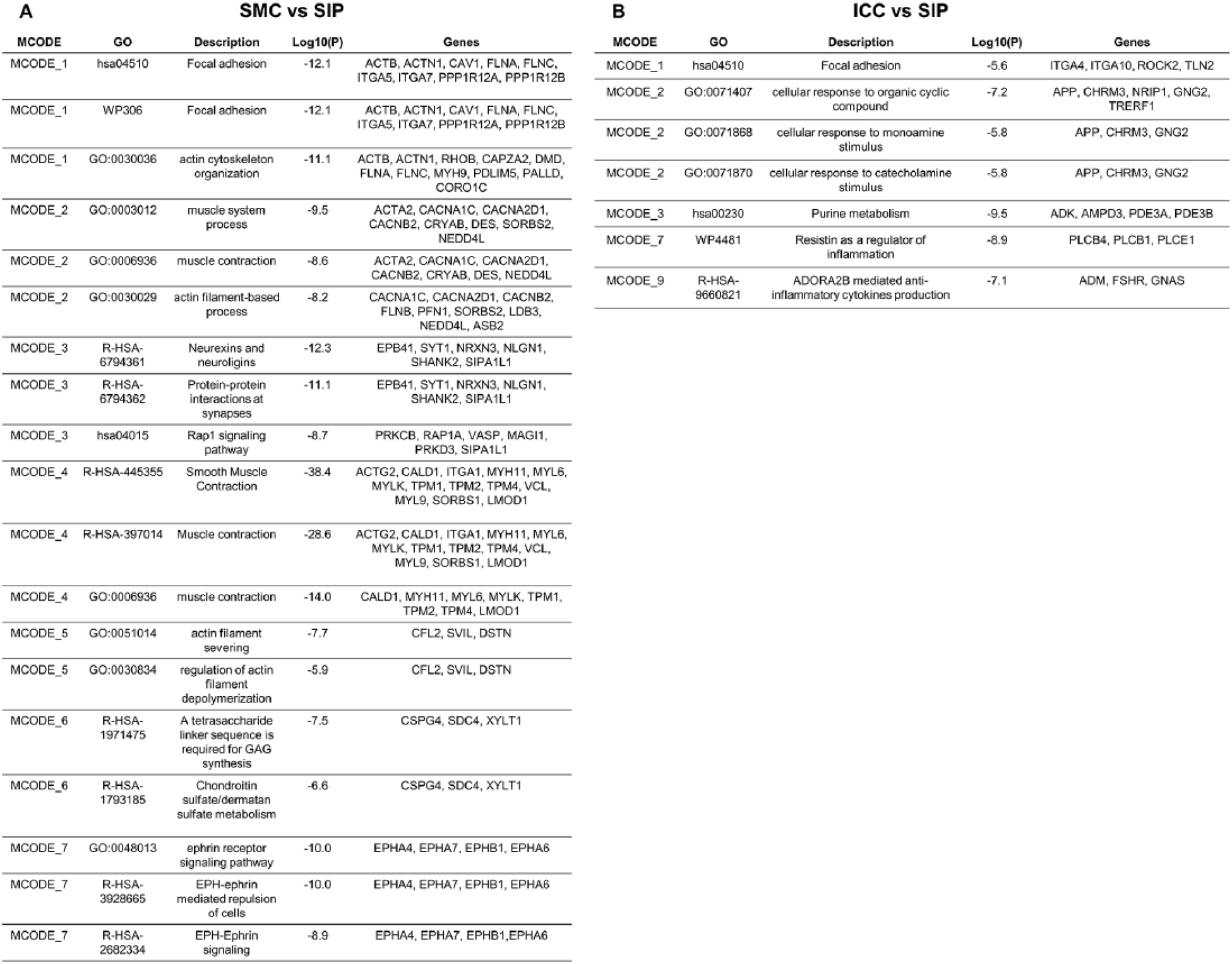
Protein-protein Interaction Enrichment Analysis - Differential gene expression of SIP syncytium cell types compared to other SIP syncytium cells. (A-B) Metascape protein-protein interaction networks composed of >3 differentially overexpressed genes for each SIP syncytium cell type were subjected to MCODE process enrichment analysis. Representative GO Terms corresponding to densely connected protein-protein network components are shown for SMCs (A) and ICCs (B). Metascape analysis did not identify networks of >3 differentially overexpressed genes for PαCs compared to other SIP syncytium cells. Differentially expressed genes were determined by comparing gene expression in each SIP syncytium cell type to the other SIP cell type clusters.

Limiting our analysis to SIP syncytium cells identified many of the same differentially expressed synaptic proteins and axon guidance molecules (Figure 14), ion channels (Figure 15), ECM proteins (Figure 16), and transcriptional regulators (Figure 17), as when all cells in the dataset were included. However, we also found many additional genes differentially expressed among SIP syncytium cells that were missed in our prior analyses (Figures 7-10). For example, 48% of the synaptic proteins and axon guidance molecules differentially expressed between SIP syncytium cells (Figure 9 versus Figure 14 gene lists) are also expressed in enteric glia as well as other non-SIP cell types and thus were not identified in our initial analysis. In contrast, most ion channels and channel-associated proteins differentially expressed between SIP syncytium cells were identified even in the analysis which includes non-SIP cell types (Figure 8 versus Figure 15 gene lists). This suggests only a few of the ion channels differentially expressed between SIP syncytium cells were abundantly expressed in non-SIP cell types (7/51 (13%); Figure 15). Another interesting observation is that PαCs differentially express at high levels many more ECM components and ECM remodeling proteins compared to SMCs or ICCs (Figure 16). Collectively, these observations highlight many genes differentially expressed among SIP syncytium cells that have not yet been well-studied in these cell populations.

**Figure 14.**
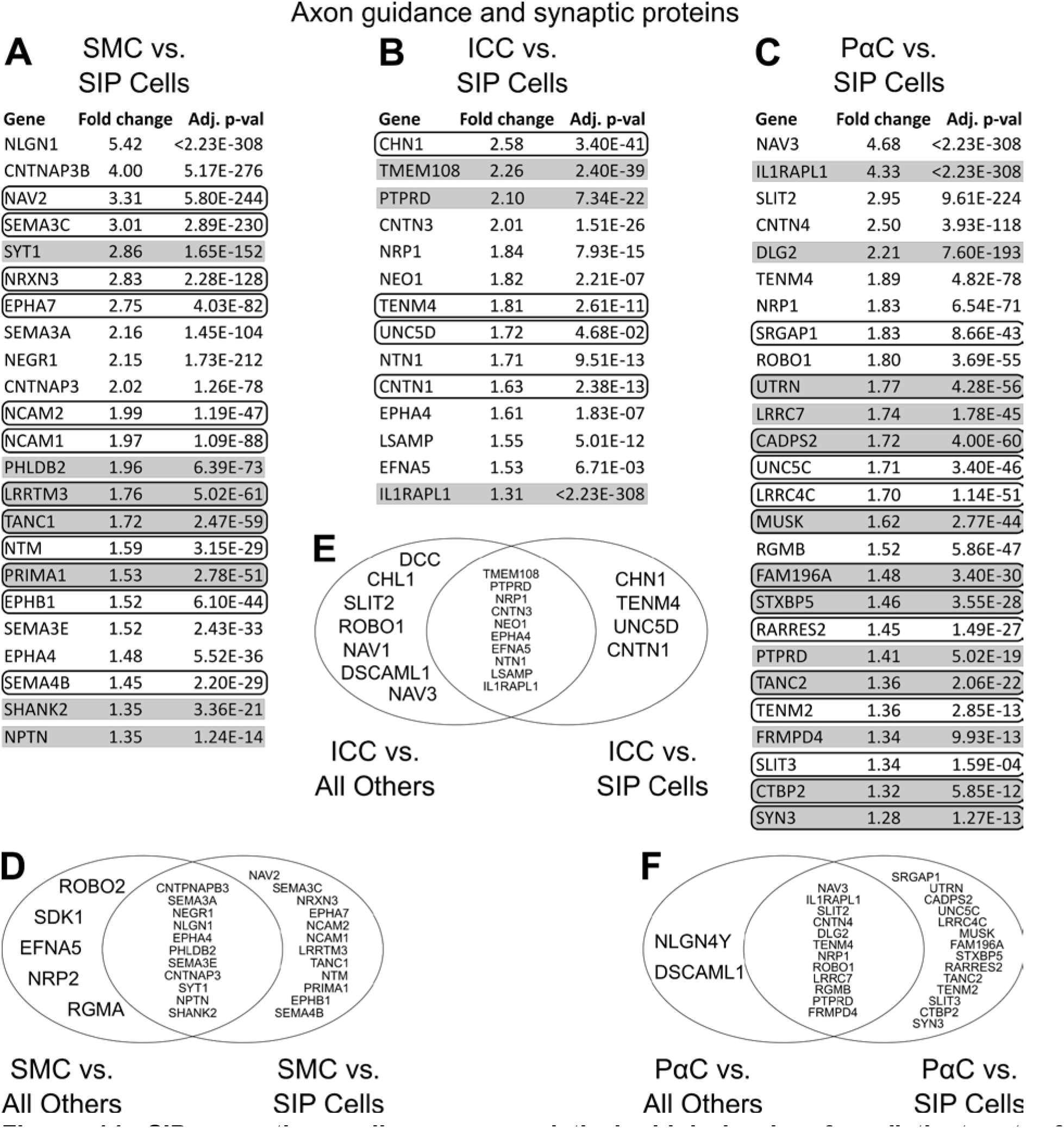
SIP syncytium cells express relatively high levels of a distinct set of transcripts encoding axon guidance and synapse function/maintenance proteins. (A-C) Genes encoding synapse-associated proteins identified as more abundant in (A) SMCs, ICCs, or (C) PαCs when these cells are compared to other SIP syncytium cell types. Genes encoding synapse-associated proteins are highlighted in gray. Genes that were not identified as differentially expressed when SIP syncytium cells were compared to all other cells in the dataset are circled. Fold change = mean number of transcripts in cluster of choice/mean number of transcripts in the other two SIP syncytium clusters. (D, E, F) Venn diagrams compare synapse-associated genes identified as enriched in SMCs (D), ICCs (E) or PαCs when these cell types are compared to all other cells in the dataset verses genes identified as enriched when these cell types are compared only to other SIP syncytium cell types.

**Figure 15.**
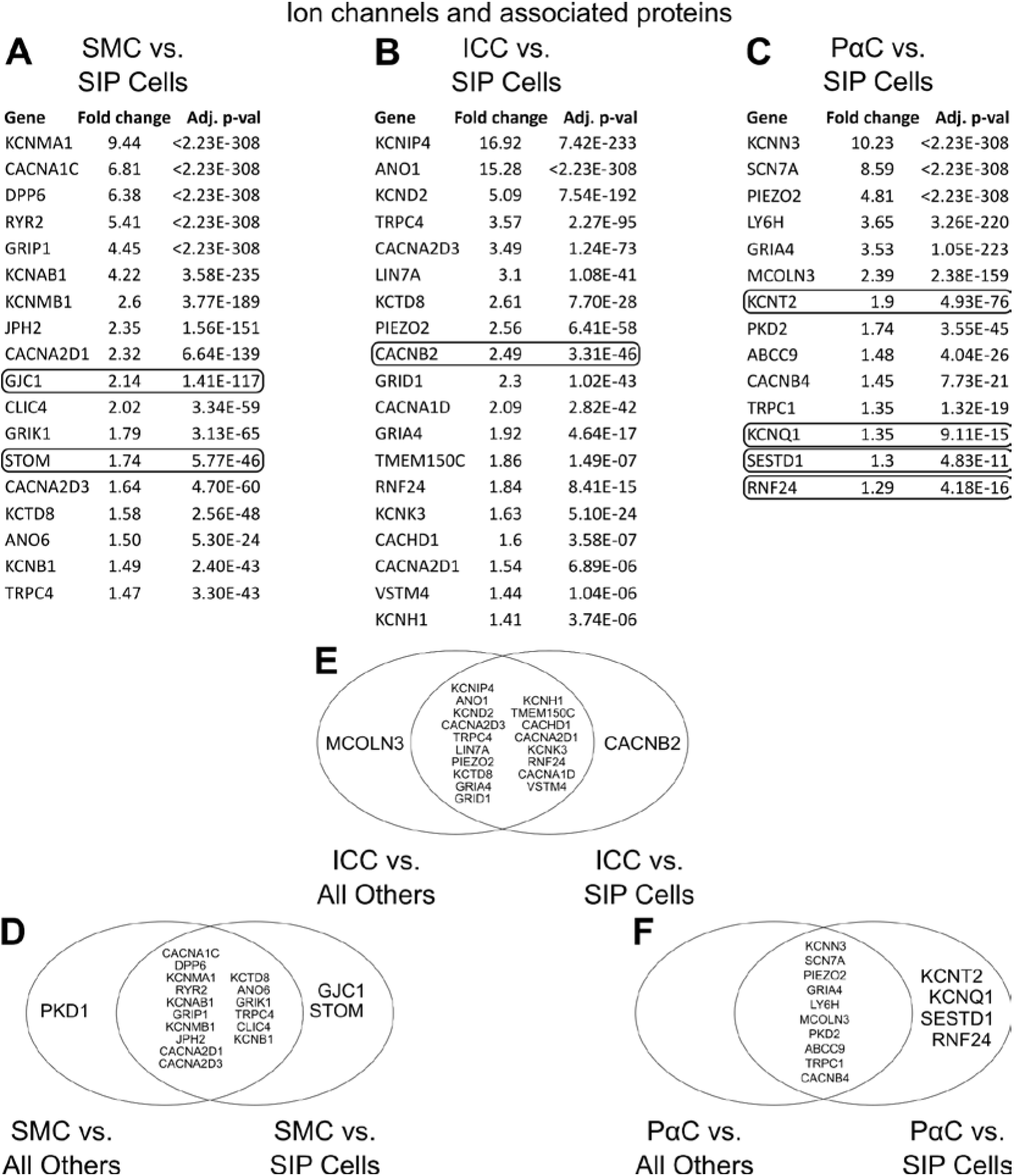
SIP syncytium cells have upregulated expression of a distinct set of transcripts encoding ion channels and ion channel-associated proteins. (A-C) Ion channels and ion channel-associated genes identified as more abundant in (A) SMCs, (B) ICCs, or (C) PαCs compared to other SIP syncytium cell types. Genes that were not identified as differentially expressed when SIP syncytium cells were compared to all other cells in the dataset are circled. Fold change = mean number of transcripts in cluster of choice/mean number of transcripts in the other two SIP syncytium clusters. (D-F) Venn diagrams compare ion channel-associated genes identified as enriched in SMCs (D), ICCs (E) or PαCs when these cell types are compared to all other cells in the dataset verses genes identified as enriched when these cell types are compared only to other SIP syncytium cell types.

**Figure 16.**
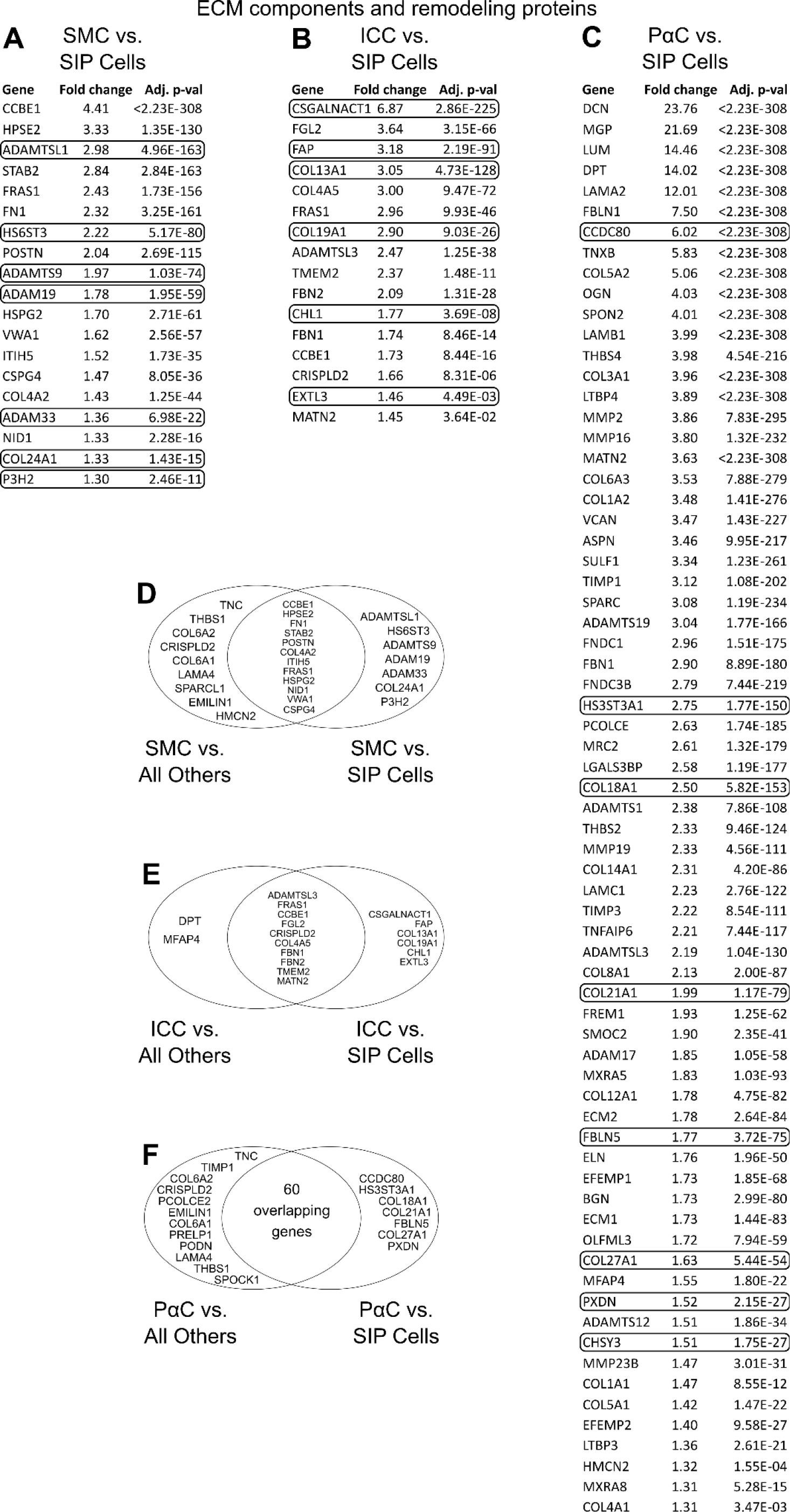
SIP syncytium cells have distinct gene expression patterns for ECM components and ECM remodelers. (A-C) Structural and nonstructural ECM components and ECM remodelers identified as more abundant in (A) SMCs, (B) ICCs, or (C) PαCs when these cells are compared to other SIP syncytium cell types. Genes that were not identified as differentially expressed when SIP syncytium cells were compared to all other cells in the dataset are circled. Fold change = mean number of transcripts in cluster of choice/mean number of transcripts in the other two SIP syncytium clusters. (D-F) Venn diagrams compare ECM-associated genes identified as enriched in SMCs (D), ICCs (E) or PαCs when these cell types are compared to all other cells in the dataset verses genes identified as enriched when these cell types are compared only to other SIP syncytium cell types.

**Figure 17.**
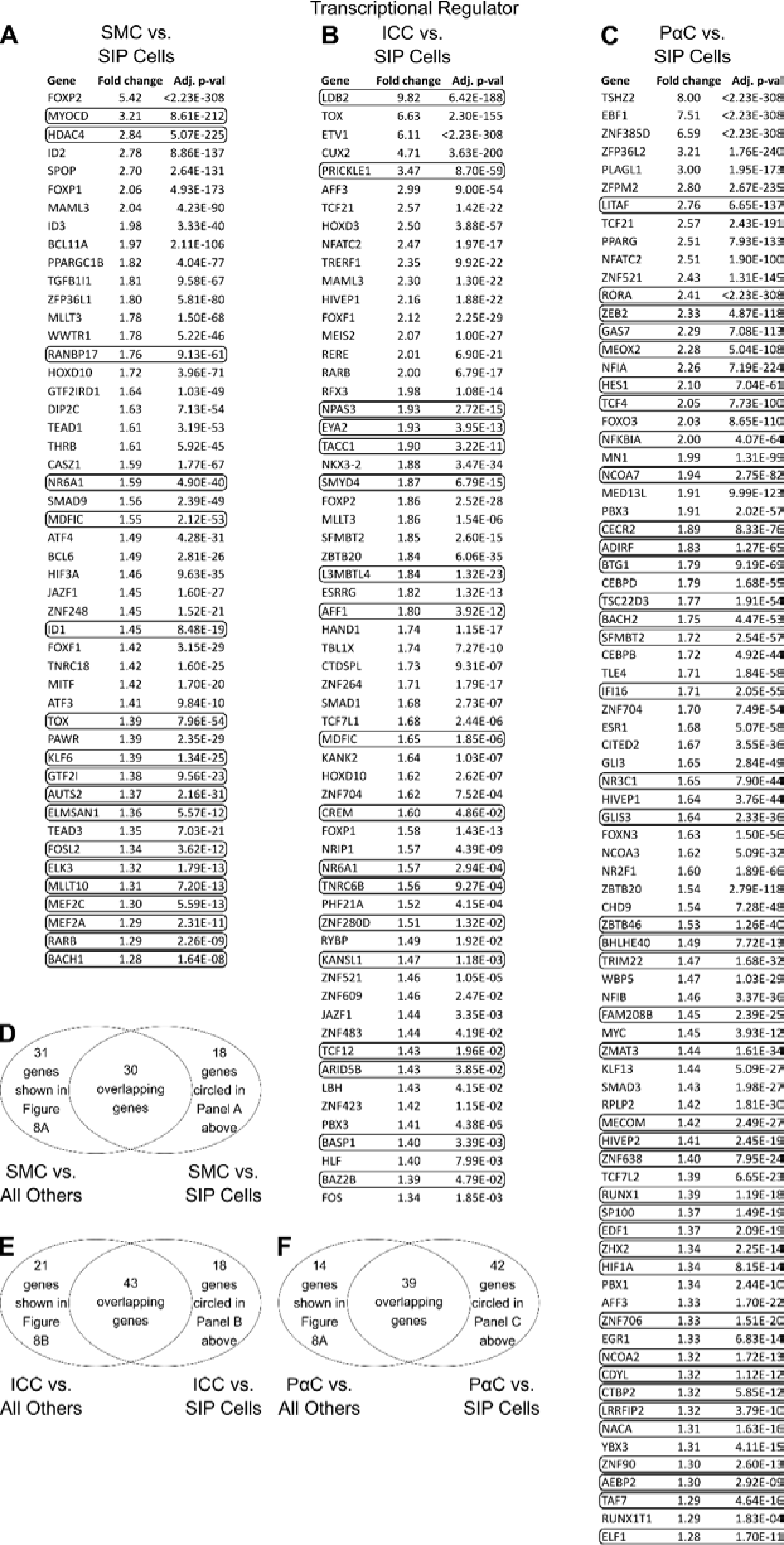
SIP syncytium cells have distinct gene expression patterns for transcriptional regulators. (A-C) Transcriptional regulators identified as more abundant in (A) SMCs, (B) ICCs, or (C) PαCs when these cells are compared to other SIP syncytium cell types. Genes that were not identified as differentially expressed when SIP syncytium cells were compared to all other cells in the dataset are circled. Fold change = mean number of transcripts in cluster of choice/mean number of transcripts in the other two SIP syncytium clusters. (D-F) Venn diagrams compare transcriptional regulators identified as enriched in SMCs (D), ICCs (E) or PαCs when these cell types are compared to all other cells in the dataset verses genes identified as enriched when these cell types are compared only to other SIP syncytium cell types.

### SIP syncytium cell types express a variety of growth factor and neurotransmitter receptors

SIP syncytium cells express at relatively high levels many cell surface receptors and ligands (Figure 18). These include tyrosine kinase growth factor receptors like *FGFR2* and *IGF1R* that are preferentially expressed in SMCs, and receptors *FGFR1*, *NTRK2*, and *KIT* that are preferentially expressed in ICCs. PαCs preferentially express *PDGFRA* (by definition) as well as *FGFR1*, *GHR*, and *CNTFR*. In addition, SIP syncytium cells differentially express cell surface receptors that regulate differentiation (TGFß receptors, BMP receptors, Hedgehog receptor *PTCHD1*) and receptors for neurotransmitters. In particular, vasoactive intestinal peptide receptor *VIPR2*, calcitonin receptor *CALCRL*, and adrenergic receptor *ADRA1A* are preferentially expressed in PαCs, whereas acetylcholine receptors *CHRM2* and *CHRM3*, and calcitonin gene related peptide (CGRP) receptor *RAMP1*, are preferentially expressed in SMCs. Interestingly, PαCs express *KITLG*, a trophic factor that activates KIT to support survival of ICCs. Ligands for TGF receptors, FGF receptors, and PDGF receptors are produced in many SIP syncytium cells and might even work cell-autologously in some cases. These observations provide many avenues for additional investigation.

**Figure 18:**
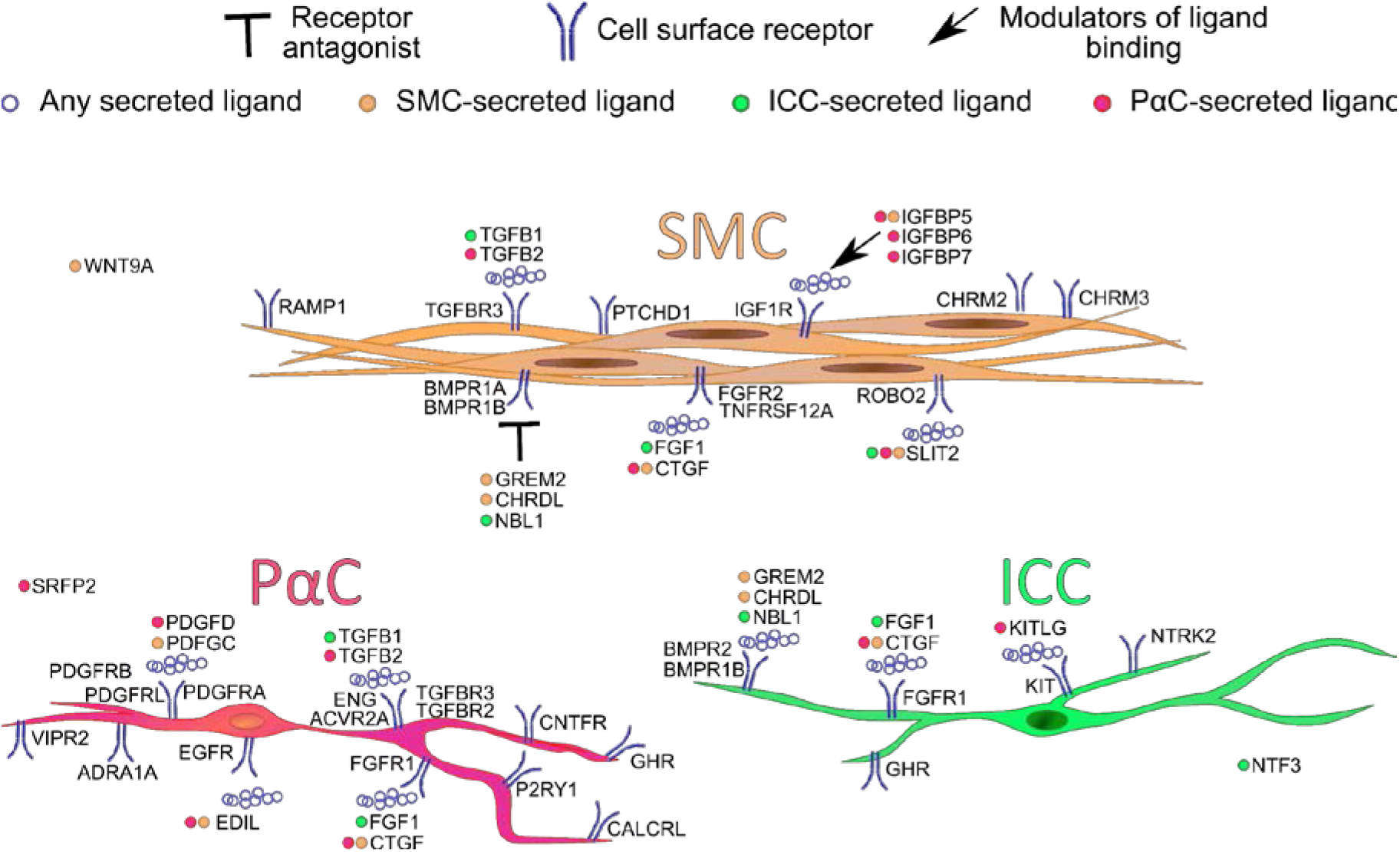
Cell Surface Receptors and Receptor-Ligand Pairs in the SIP syncytium. Many cell surface receptors for trophic factors, morphogens, axon guidance molecules, and their corresponding ligands or soluble signaling modulators are expressed by SIP syncytium cells. Neurotransmitter receptors *RAMP1*, *CALCRL*, *VIPR2*, *CHRM2*, *CHRM3*, *P2RY1*, and *ADRA1A* are also differentially expressed at relatively high levels among SIP syncytium cells. These patterns suggest interesting cross-talk between SIP syncytium cells. In some cases ligands and corresponding receptors are produced in the same cell. For other receptors, ligands are expressed by surrounding cell types that may not be included in this image. Colored dots adjacent to ligand symbols correspond to cell type of origin.

### The two PαC populations differ in expression of ion channels and structural ECM mRNA

Given previous reports of distinct PαC subtypes^17^ and two PαC clusters identified in our UMAP projections (Figure 2C), we compared gene expression in PαC#1 to PαC#2 (Figure 19A). *PDGFRA* expression was significantly increased in both PαC clusters compared to all other cells in the dataset (Figure 3G). While both PαC clusters (PαC#1 and PαC#2) expressed *KCNN3* (Figure 3H) and *P2RY1* (Figure 3I) at higher proportions than any other cell type, these transcripts were much more abundant in PαC#1 than in PαC#2 and only significantly different in PαC#1 when compared to all other clusters. Also, compared to PαC#2, the PαC#1 cluster had higher levels of the mechanosensitive ion channel *PIEZO2*, neurotransmitter receptors *ADRA1A* and *RAMP1* (Figure 19A), and a greater abundance of transcripts encoding axon guidance and synaptic proteins (Figure 19B and Figure 20A, B), ion channels and ion channel-associated proteins (Figure 19C and Figure 20C, D), and transcriptional regulators (Figure 19D and Figure 20E, F). In contrast, PαC#2 express a greater number of structural ECM components (Figure 19E-G and Figure 20G, H).

**Figure 19.**
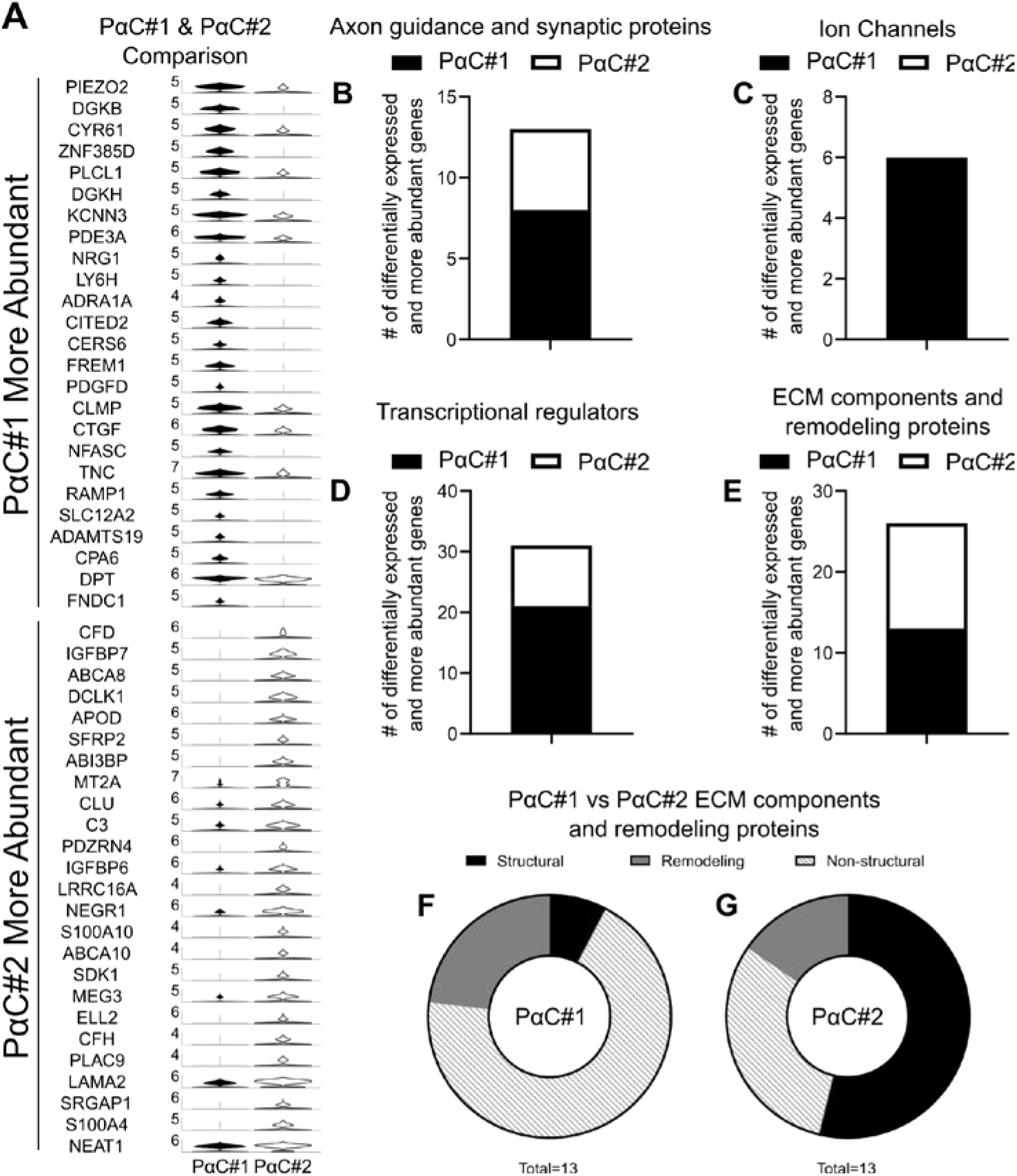
Two PαC clusters have distinct gene expression profiles. (A) Violin Plots of the 50 most differentially expressed genes between clusters PαC#1 and PαC#2 show the distribution of gene expression levels for individual nuclei. Expression level is defined as log_e_(mean number of transcripts in PαC cluster of choice/mean number of transcripts in other PαC cluster). (B) Clusters PαC#1 and PαC#2 express similar numbers of genes involved in synapse function or maintenance and axon guidance. (C) Ion channel and ion channel-associated genes were more abundantly expressed in PαC#1 compared to PαC#2. (D) A greater number of transcriptional regulators are more abundantly expressed in PαC#1 compared to PαC#2. (E-G) Cells in clusters PαC#1 and PαC#2 express similar numbers of ECM components and ECM remodeler mRNA (E), however, PαC#2 express a greater proportion of structural ECM components compared to PαC#1 (F-G). Numbers below pie charts indicate the number of genes represented in the pie chart.

**Figure 20.**
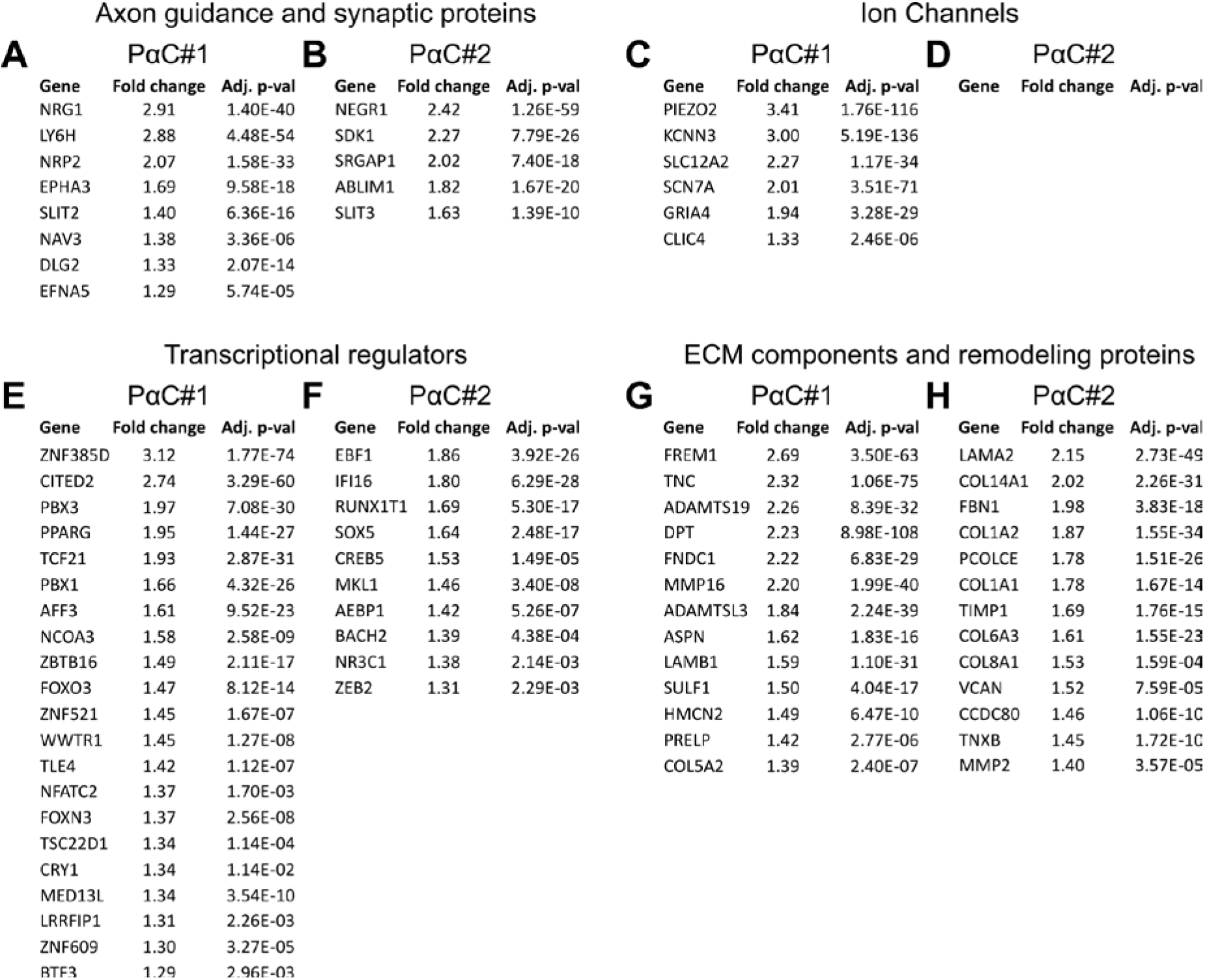
Two distinct PαC clusters differentially express axon guidance and synaptic proteins, ion channels, transcriptional regulators, ECM components and ECM remodelers. (A-D) Relative abundance of differentially expressed genes in PαC#1 or PαC#2 when these cell type clusters are compared to each other. (A, B) Synapse function or maintenance and axon guidance genes enriched in PαC#1 (A) or PαC#2 (B). (C, D) Ion channels and ion channel-associated genes enriched in PαC#1 (C) or PαC#2 (D). (E, F) Transcriptional regulators enriched in PαC#1 (E) or PαC#2 (F). (G, H) ECM components and remodeling genes enriched in PαC#1 (G) of PαC#2 (H). (A-H) Adj. p-val = Bonferroni adjusted p-value for each gene. Fold Change = mean number of transcripts in PαC cluster of choice/mean number of transcripts in the other PαC cluster.

### The PαC populations are distinct from fibroblasts

Since PαCs were previously described as fibroblast-like cells^26^, we compared the two PαC clusters against the other fibroblast cluster in our dataset (identified by *MEOX2*, *COL1A2*, *COL1A1*, *FMO1*, *LSP1,* and *VIM* expression). Metascape network analysis showed that most of the GO term enrichment for genes involved in nervous system development came from PαC#1 (Figure 4E). Clusters PαC#1 and PαC#2 had few differentially expressed genes in common, whereas PαC#2 and the fibroblast cluster shared a larger proportion of differentially expressed genes. PαC#1 and fibroblasts had no differentially expressed genes in common (Figure 4F). Protein-protein MCODE Interaction Enrichment Analysis (PPMIEA), which provides GO terms for differentially expressed genes when >3 encoded proteins interact with each other, showed that the most enriched interaction networks for PαC#1 involved ribosomal proteins (Figure 21A). Other enriched networks for PαC#1 included smooth muscle contraction and muscle development, purinergic and nitrergic signaling, and anti-inflammatory cytokine production. In contrast, while PαC#2 are enriched for the complement system in neuronal development and plasticity, most other enriched protein-protein interaction networks were ECM-related (Figure 21B). The fibroblast cluster also primarily showed enrichment of ECM-related networks, similar to PαC#2 (Figure 21C).

**Figure 21:**
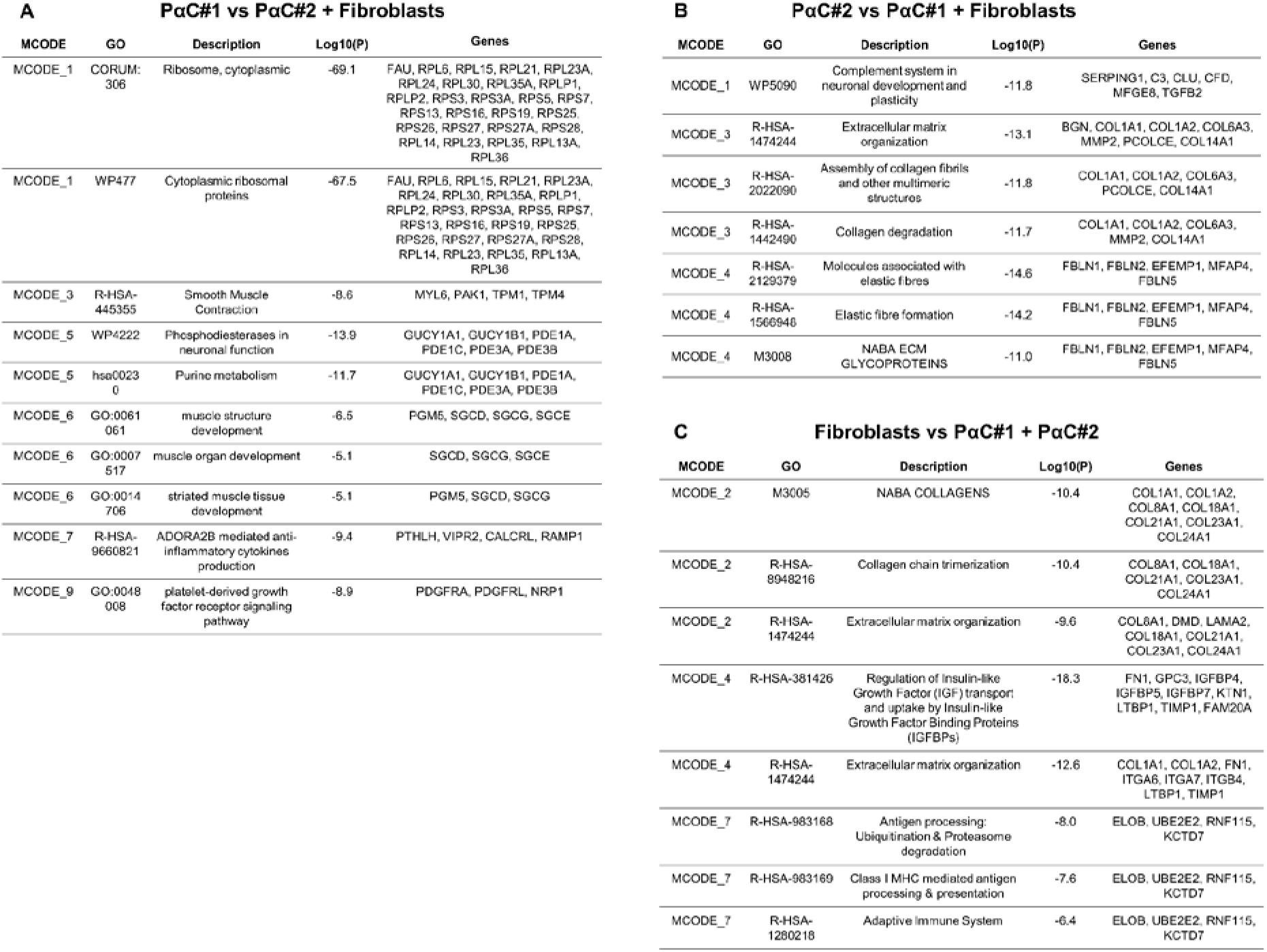
Protein-protein Interaction Enrichment Analysis comparing gene expression of the two PαC clusters versus *VIM*/*MEOX2*/*FMO1*/*LSP1*-expressing fibroblasts in the dataset. (A-C) Metascape protein-protein interaction networks composed of >3 differentially expressed genes found at elevated levels in each of the three fibroblast-like cell types (PαC#1, PαC#2, or fibroblasts) compared each other were subjected to MCODE process enrichment analysis. Representative GO Terms correspond to densely connected protein-protein network components for nuclei in PαC#1 cluster (A), PαC#2 cluster (B), and fibroblasts (C).

### Right colon has greater abundance of transcripts for ion channels and transcriptional regulators, while left colon expresses more transcripts for axon guidance and synaptic proteins

Since the colon has regional differences in motility and epithelial cell composition^27^, we wondered if gene expression of SIP syncytium cells differs between right (ascending) and left (sigmoid) colon. We found nuclei of SIP syncytium cells organized neatly into 3 distinct clusters (Figure 22A, B). Forced comparison between right and left colon showed that SIP cell types had similar mean UMIs and mean numbers of genes detected in each bowel region (Figure 22C). A majority of nuclei corresponding to PαC#1 came from the right colon, while right and left colon contributed similar number of PαC#2 to our dataset (Figure 22D). The most differentially expressed genes between the right and left colon are shown in Figure 22E-G. The number of significantly differentially expressed genes was limited in ICCs, perhaps because of relatively low cell numbers (right colon: 373 nuclei, left colon: 207 nuclei), (Figure 22G).

**Figure 22.**
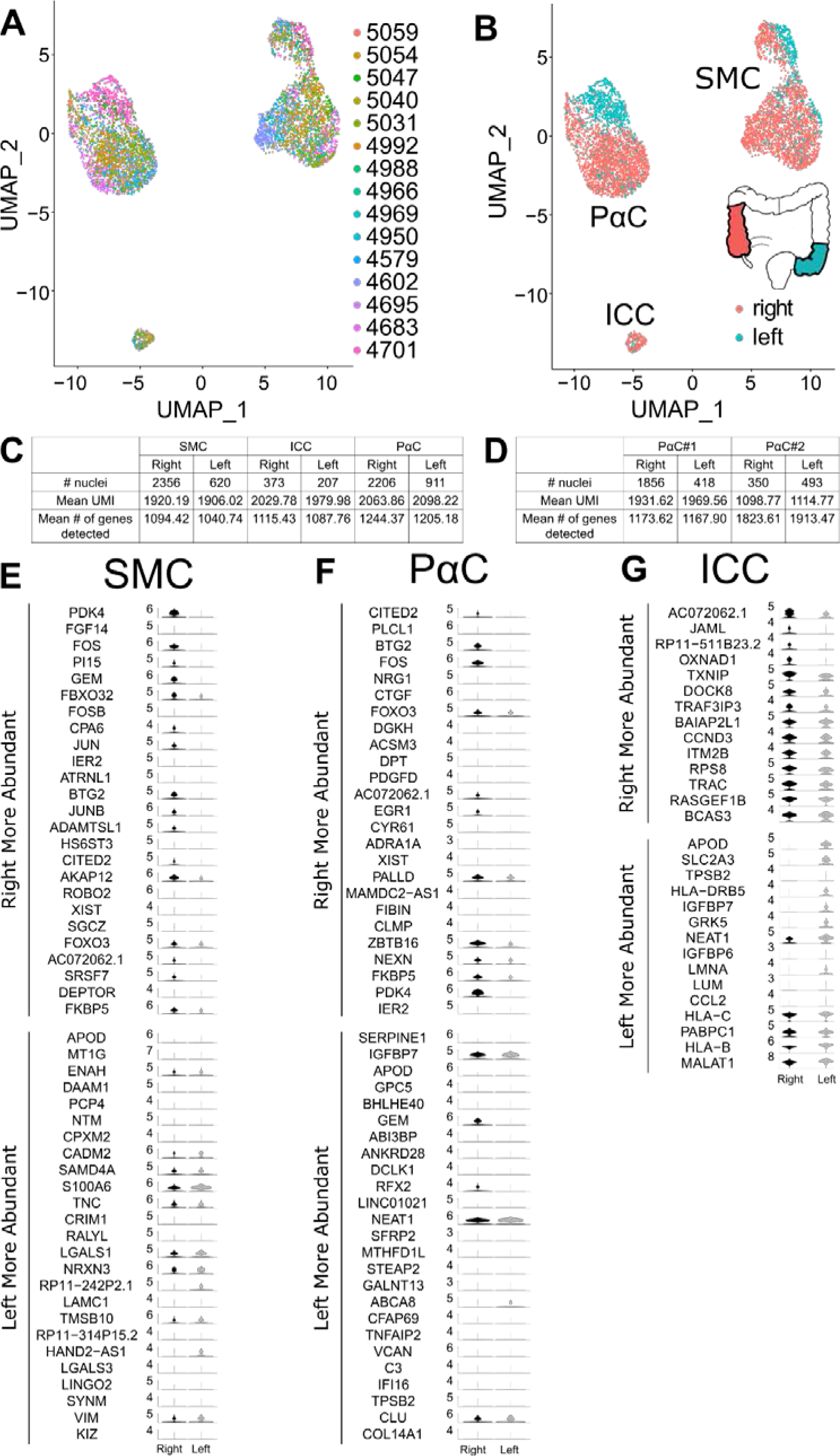
Gene expression in SIP syncytium cells differs in right versus left colon. (A) SIP syncytium nuclei cluster by cell type but not by sample ID after removing non-SIP cell types from the dataset. Nuclei are color coded by sample ID. (B) Color coding shows partial segregation of gene expression by bowel region of origin. (C) Nuclei from the left or right colon express similar numbers of unique RNA molecules (UMI) and unique genes. (D) PαC#1 and PαC#2 nuclei in the left and right colon express similar numbers of UMIs and unique genes. PαC#1 nuclei were 5.3-fold more abundant in right colon samples than PαC#2 nuclei, whereas left colon had similar numbers of PαC#1 and PαC#2 nuclei. (E-G) Violin Plots of the 50 most differentially expressed genes in right/ascending colon versus left/sigmoid colon for SMC (E), PαC (F), and ICC (G) show the distribution of gene expression levels per individual nucleus. Expression level is defined as log_e_(expression level in nucleus/mean expression level for nuclei across all clusters).

In contrast, compared to left colon, right colon PαCs (Figure 23A, B) and SMCs (Figure 24A, B) had more transcripts encoding axon guidance and synaptic function/maintenance proteins, many more ion channels and ion channel-associated proteins (**PαC**: Figure 23C, D**; SMC:** Figure 24C, D), and more transcriptional regulators (**PαC**: Figure 23E, F**; SMC:** Figure 24G, H). Only a few neurotransmitter receptors were found to be differentially expressed between the two bowel regions, and transcripts for most of these were significantly more abundant in the right colon. *ADRA1A* and *CALCRL* were more abundant in right colon PαCs, and the two muscarinic acetylcholine receptors *CHRM2* and *CHRM3* were more abundant in right colon SMCs. Only the tachykinin receptor *TACR2* is more abundant in left colon SMCs. While right and left colon PαCs and SMCs expressed a similar number of ECM components and extracellular matrix remodeling proteins (**PαC**: Figure 23G, H**; SMC:** Figure 24E, F), right colon PαCs expressed a lower number of structural ECM components compared to the left colon (Figure 23I, J). Collectively, these data identify regional differences in SMC and PαC gene expression that might be linked to differences in bowel motility.

**Figure 23:**
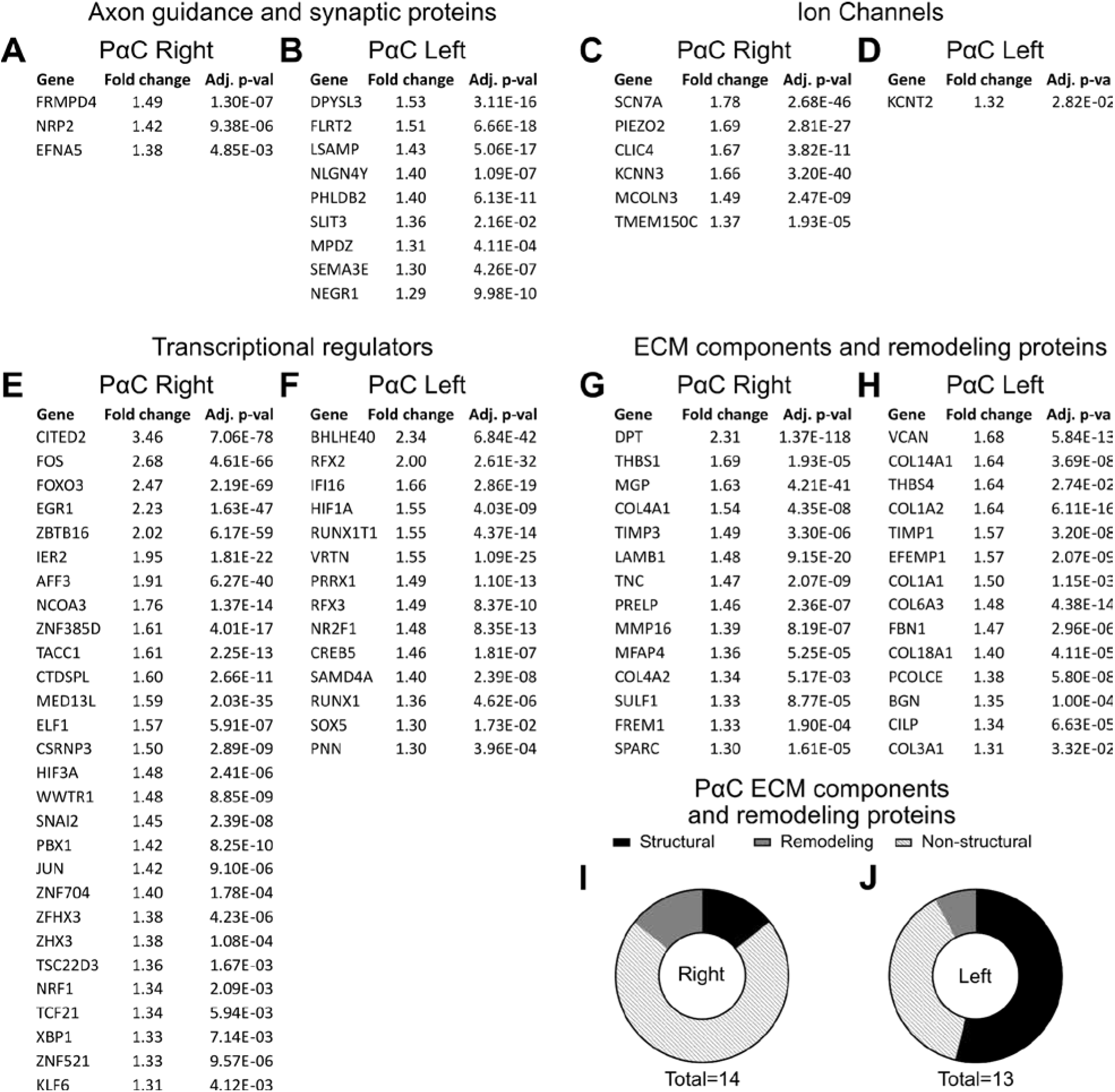
PαC from the right and left colon have unique gene expression patterns. (A-H) Differentially expressed genes that are more abundant in either right or left colon PαC. (A, B) Axon guidance and synaptic genes. (C, D) Ion channels and ion channel-associated genes. (E, F) Transcriptional regulators. (G, H) ECM components and remodeling genes. (A-H) Adj. p-val = Bonferroni adjusted p-value is shown for each gene. Fold Change = mean number of transcripts in PαC cluster of choice/mean number of transcripts in other PαC cluster. (I, J) Pie charts show different gene expression patterns for ECM structural, non-structural, and remodeling genes in left and right colon-derived PαC clusters. Numbers below pie charts indicate the number of genes represented in the pie chart.

**Figure 24:**
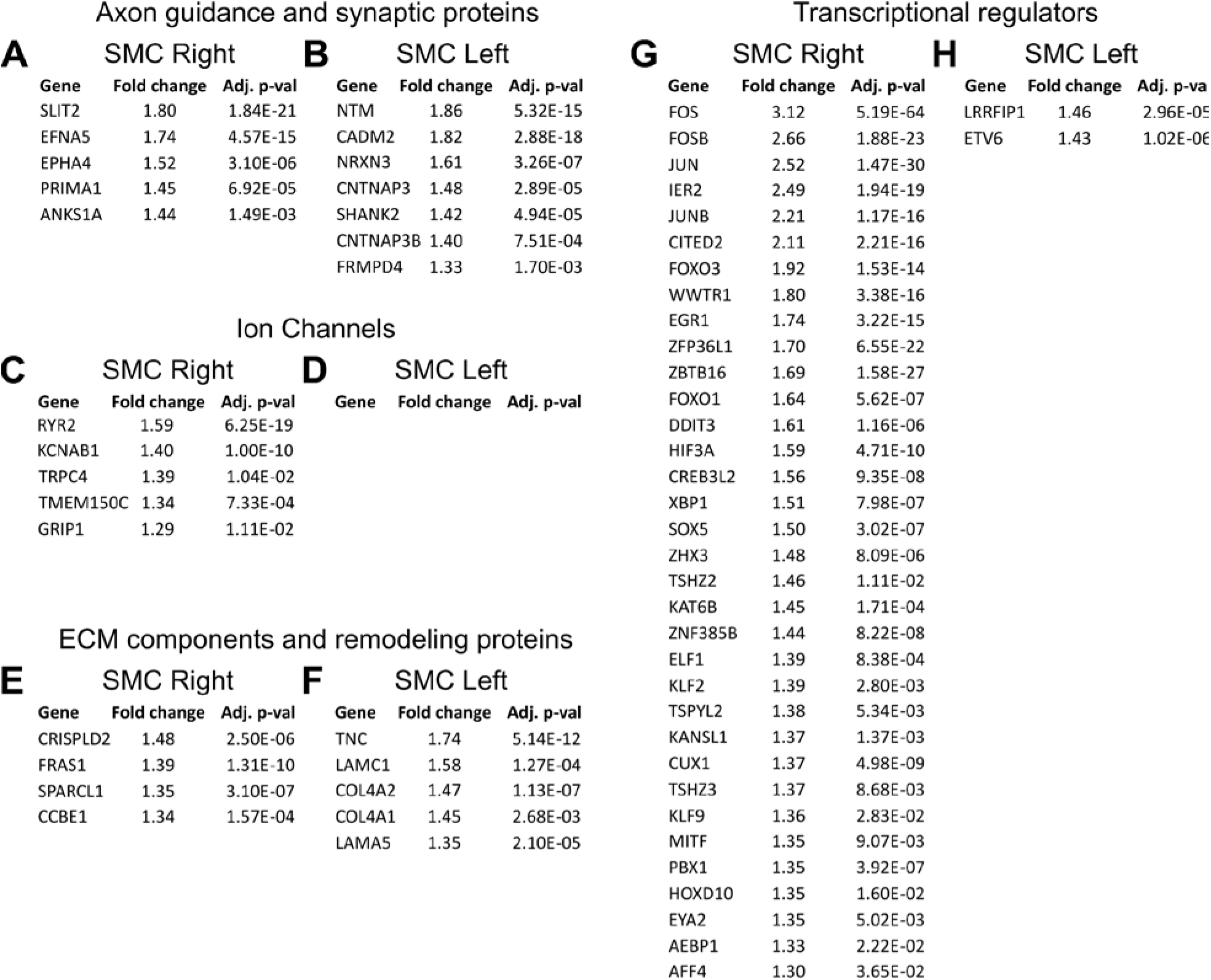
SMC from the right colon and left colon have unique gene expression profiles. (A-H) Differentially expressed genes that are more abundant in either right or left colon derived SMCs. (A, B) Axon guidance and synaptic genes. (C, D) Ion channels and ion channel-associated genes. (E, F) ECM structural, non-structural, and remodeling genes. (G, H) Transcriptional regulators. (A-H) Adj. p-val = Bonferroni adjusted p-value is shown for each gene. Fold Change = mean number of transcripts in SMC cluster of choice/mean number of transcripts in the other SMC cluster.

## Discussion

Here we present the first analysis of adult human colon single nucleus RNA sequencing focused on the SIP syncytium. Our analyses highlight novel roles and interactions for SIP syncytium cells. For example, adult SMCs express many mRNA encoding repulsive axon guidance molecules likely to restrict axon entry into the muscle layer to specific classes of enteric nervous system neurons (e.g., excitatory and inhibitory motor neurons). This is largely unexplored biology. ICCs and PαCs express mechanosensitive ion channels suggesting they directly respond to mechanical forces so prevalent in the bowel. PαCs were split into two distinct populations. PαC#1 expresses numerous ion channels consistent with the PαC role in inhibitory neurotransmission within the SIP syncytium, while PαC#2 prominently expresses structural ECM genes. In contrast, PαC#1 expresses many non-structural ECM genes and many ECM remodeling proteins. Furthermore, SMCs and PαCs in the right colon express many more transcriptional regulators and ion channels than SMCs and PαCs in the left colon, suggesting regional differences in SIP syncytium function. Finally, all SIP syncytium cell types share differentially enriched expression of at least 6 transcription factors not previously reported in this context. These proteins may act as a transcriptional network to direct SIP syncytium cell type differentiation and/or maintenance. These major findings reinforce and extend existing literature on the SIP syncytium.

Our initial analysis strategy resulted in two SMC clusters, one ICC cluster, and one PαC cluster. However, eliminating one colon sample (#5035) condensed the SMC into a single cluster, and split PαC into two clusters. One concern is that #5035 was from a young adult with sigmoid volvulus, an uncommon condition that typically occurs when the sigmoid colon is massively dilated. Massive dilation may occur because of distal obstruction or because of dysfunction in the dilated bowel and we do not know if this tissue was ischemic. We therefore omitted #5035 since our goal was to define normal SIP syncytium biology. Interestingly, the two human PαC clusters delineated after removing #5035 may correspond to the two murine PαC subpopulations distinguished by high or low PDGFRA expression (PDGFRA^high^ versus PDGFRA^low^)^17^. In mice, PDGFRA^low^ cells are marked by relatively high expression of *Cacna1g*^17^. Unfortunately we did not detect *CACNA1G* in the human PαC clusters, possibly due to the limited read depth or to differences between human and mouse. For example, our human data shows *THBS4* at relatively high levels in PαC (3.16-fold higher than all other cells) but *Thbs4* was recently described as an ICC-specific marker in mice^15^. One interesting physiologic distinction in mice is that in the setting of partial intestinal obstruction where the muscle layer increases in size, PDGFRA^low^ cells proliferate whereas PDGFRA^high^ cells hypertrophy.

Our human data show that fewer PαC#2 cluster nuclei had detectable *PDGFRA* compared to nuclei in PαC#1. This suggests PαC#2 may correspond to PDGFRA^low^ in mice, but this needs to be validated by more extensive comparisons of gene and protein expression profiles in murine^28^ and human cells. Nonetheless, the divergent gene expression profiles in PαC#1 versus PαC#2 are interesting in light of our limited understanding of PαC function. In particular, PαCs have been called “fibroblast-like cells”^26^ and also have roles in inhibitory neurotransmission within the SIP syncytium^1, 2^. Our data suggest that these attributes might be distinct functions of PαC#2 and PαC#1 respectively, since PαC#2 express higher levels of many structural ECM components, and PαC#1 express higher levels of many ion channels, channel associated proteins and neurotransmitter receptors. PαC#1 thus appears to be the more electrically and transcriptionally active cell population. If human PαC#2 are similar to PDGFRA^low^ in mice, we would expect an increase in ECM-producing PαC#2 in the setting of partial obstruction-induced muscle hypertrophy with less expansion of PαC#1, a hypothesis that could be tested if appropriate human specimens were obtained for single nucleus RNAseq analysis.

### New insights

From the functional perspective, we were not surprised to see prominent expression of many ion channels, ion channel-associated proteins, axon guidance molecules, and synaptic proteins, since all SIP syncytium cells are innervated by neurons and are electrically active^1, 3, 29^. We were surprised, however, to see mechanosensitive ion channels (*PIEZO2* and *TMEM150C*) expressed in ICCs and PαCs. This suggests that ICCs and PαC can directly detect mechanical force (e.g., from bowel distension) and possibly modulate smooth muscle contraction without requiring input from primary sensory neurons of the ENS. Our data also show for the first time (to the best of our knowledge) high level expression of vasoactive intestinal peptide receptor 2 (*VIPR2*) in the PαC cluster (Figure 15), which suggests VIP could directly modulate human PαC function. In addition, we detected relatively high levels of the CGRP co-receptors *CALCRL* in PαCs and *RAMP1* in SMCs. β-CGRP, a neurotransmitter expressed by enteric neuron subtypes like intrinsic primary afferent neurons^30, 31^, has been implicated in the intestinal peristaltic reflex in mice^32, 33^. Expression of CGRP receptors in SMCs and PαCs may explain why CGRP induces smooth muscle relaxation even when neurotransmission is blocked with tetrodotoxin^34–36^. This may have clinical relevance since recently-approved anti-CGRP migraine therapeutics such as Erenumab and Fremanezumab can cause prominent gastrointestinal symptoms including constipation, diarrhea, abdominal pain, and nausea^37–39^. Our data suggests that SIP cells may respond to neurotransmitters in ways that remain to be delineated by functional experiments. In addition to neurotransmitter receptors and ligands, the expression of genes for cell surface receptors and ligands for several growth factors, differentiation factors, and axon guidance molecules suggests significant functional chemical crosstalk between the SIP syncytium cell types.

Another interesting aspect of SIP syncytium cell biology is that murine ICC and SMC share a common embryonic KIT+ precursor at least in the longitudinal smooth muscle layer^40–42^. This common precursor also expresses PDGFRα and PDGFRß receptors and during fetal development, the ligands PDGF-A and PDGF-B are expressed in circular muscle SMC and in enteric neurons, respectively. Blocking PDGFR with the chemical antagonist AG1295 suppresses SMC differentiation in the longitudinal muscle layer and appears to induce formation of ICC^43^. The origin of mature PαC remains unclear, but it is tempting to speculate that PDGFR-expressing mesenchymal cells are a common precursor for all three SIP syncytium cell types at least during fetal development^17, 43^. However, this has not yet been experimentally confirmed. The wide-spread expression of PDGFRA in mesodermal and ectodermal derivatives during embryonic organogenesis^44^ and in the adult mouse, where PDGFRA is expressed in most but not all fibroblast populations^45^, means that we will need to delineate functional differences in bowel PαC subpopulations based on additional cell type-specific markers.

Supporting their close developmental relationship, all SIP syncytium cell types express high levels of six transcriptional regulators relative to other cells in our dataset (*MEIS1*, *MEIS2, PBX1, FOS, ZBTB16,* and *SCMH1*) (Figure 11B). Finding MEIS1, MEIS2, and PBX1 in the same cells is not surprising since MEIS1 and MEIS2 orthologs regulate PBX1 translocation to the nucleus and MEIS1 and PBX1 heterodimerize on DNA to control gene expression, including for a set of HOX proteins^46^. The immediate early gene *FOS* mediates TGF-β signaling and TGF-β receptors are expressed by all SIP syncytium cell types in our dataset (Figure 18). TGF-β signaling has also been shown to be important for differentiation of ICC and vascular smooth muscle^47, 48^. *ZBTB16/PLZF* regulates cellular responsiveness to FGF signaling^49^ and was recently described as a likely causative genomic locus in a rat model of myocardial hypertrophy, fibrosis, and hypertension^50^. FGF receptors are expressed by all SIP syncytium cell types in our dataset (Figure 18). *SCMH1* is a component of the polycomb repressive complex 1 that regulates expression of large numbers of genes through chromatin modifications^51^. These differentially expressed factors may be part of a shared SIP-specific transcriptional regulatory network. Supporting this hypothesis, when the gene expression patterns of only SIP syncytium cell types is compared against each other, these transcriptional regulators are either no longer detected as differentially expressed (*MEIS1*, *SCMH1*, and *ZBTB16*) or only upregulated in a single SIP syncytium cell type (*FOS*, *MEIS2*, and *PBX1*).

A particularly curious observation is that SMCs and PαCs in the ascending/right colon express more transcriptional regulators, ion channels (Figures 22-25), and neurotransmitter receptors than these same cell types in sigmoid/left colon. Interestingly, this correlates with 5.3-fold more PαC#1 compared to PαC#2 nuclei in our right colon data, compared to a ratio of 0.85 to 1 for these cells in the left colon (Figure 22D). The increased right colon transcriptional, electrical, and chemical complexity in the SIP syncytium appears to parallel the observation that ENS circuits are also more complex in proximal (right) compared to distal (left) colon^27^. In contrast, expression of a greater number of structural ECM genes in left colon PαCs compared to right colon PαCs may correlate with the comparatively thicker muscularis layer with denser ECM in the sigmoid colon. To the best of our knowledge, this is the first report of regional differences in gene expression for colon SMCs and PαCs.

After completing targeted analysis based on known or suspected roles of the SIP syncytium, we pursued systems-level analysis^23^ to identify unexpected protein networks or potential SIP syncytium functions. The Metascape protein-protein MCODE Interaction Enrichment Analysis (PPMIEA) (Figures 5 and 13) highlights interesting aspects of SIP syncytium biology. As might be expected, GO terms for smooth muscle include “Smooth Muscle Contraction” and “Muscle Contraction”, based on many proteins of the contractile apparatus. Interestingly, compared to other cells in our dataset, SMC are enriched in mRNA for focal adhesion-associated proteins, proteins that metabolize heparan sulfate and chondroitin sulfate proteoglycans, and axon guidance molecules. GO terms for ICC include “Purine metabolism”, driven in part by *GUCY1A1* and *GUCY1B1*. These soluble guanylate cyclase subunits make cGMP in response to nitric oxide, and ICC also express phosphodiesterases that degrade cGMP as previously reported^52^. In addition, ICC GO terms highlight interactions of ICC with neurons (“Neuronal System”, “Protein-Protein Interaction at Synapses”) when ICC are compared to all other cells and highlight responses to monoamines/catecholamines when ICC are compared to other SIP syncytium cells. In contrast, GO terms for PαC versus all other cells highlight expression of many extracellular matrix components (collagens, fibrillin, tenascin C, laminins, fibulins), the collagen-specific endoplasmic reticulum chaperone SERPINIH1, and TIMP1 that prevents ECM degradation, as well as proteins that permit direct responses to nitric oxide, CNTF, LIF, IL6 and OSM^53^. IL6 and OSM have potent pro-inflammatory roles in Crohn’s disease^54, 55^, where striking proliferation and hypertrophy of the bowel muscle layer causes fibrostenosing strictures^56^. This suggests that thickening of the muscularis propria in the setting of inflammation may at least in part be attributed to proliferation of PαC that are already known to proliferate in some contexts^17^.

This study has several limitations. To generate these data we microdissected cells from around the region of the myenteric plexus. These cells segregated into 14 clusters that were readily identified as SMC, PαC, ICC, B cells, T cells, lymphatic and vascular endothelial cells, fibroblasts, mast cells, macrophage, neurons, and glia based on well established cell markers. While it might be tempting to assume that the relative ratio of nuclei in this dataset reflects the ratio of microdissected cells, some cell types appear much more likely to produce evaluable RNAseq data than others with our methods. For example, our nuclear isolation strategy generated data from 48 neurons and 3798 glia (79 to 1 glia to neuron ratio)^18^ but our manual counting of ENS cells in three-dimensional Z-stacks from these same human colons indicates that there are 3 to 5.5 glial cells per neuron in the myenteric plexus region^57^. Thus, glial nuclei were ∼20-fold more likely than neuronal nuclei to produce RNAseq data. The relative likelihood of generating RNAseq data from other bowel cells is not established. Furthermore, because we isolated tissue near the myenteric plexus, we did not capture distinct types of ICC found in different layers of the bowel^16^. As with many single-nucleus sequencing studies, our observations are limited by low read depth. We observed many genes that appeared to be expressed in only a small proportion of nuclei in a given cluster; however, it is possible more cells expressed the gene, but at too low a level to be detected by our assay. The observations based on our transcriptomics data were not followed by protein-level validation of RNA expression patterns in SIP syncytium cells. However, our RNAseq data correlate well with known RNA and protein data from the SIP syncytium. Furthermore, ENS gene expression patterns in this same dataset were previously validated and correlate well with the prior ENS literature as detailed in our manuscript^18^. Despite these limitations, these analyses provide deep insight into SIP syncytium biology, revealing previously unrecognized gene expression patterns for many gene classes. We believe this first-of-its-kind analysis of human colon data will facilitate fascinating hypotheses to guide future studies of the SIP syncytium. The work may have relevance for human bowel motility disorders and for understanding pathologic effects of bowel inflammation or obstruction.

## Preprint server

### Data Transparency

Full data sets are deposited in Gene Expression Omnibus: GEO accession number GSE156905. All lists of differentially expressed genes generated in this analysis can be accessed as supplementary data files with this manuscript.

### Author Contributions

Conceptualization, S.S. and S.K.H.; Methodology, S.S. and S.K.H.; Investigation, S.S., S.K.H., A.J.T., and D.K.; Formal Analysis, A.J.T., D.K., S.S., and S.K.H.; Data Curation, A.J.T. and D.K.; Writing – Original Draft, S.S. and S.K.H.; Writing – Review & Editing, S.S., S.K.H,. A.J.T., D.K., C.M.W., R.O.H.; Resources, S.S., C.M.W., and R.O.H; Supervision, S.S. and S.K.H; Funding Acquisition, R.O.H.

## Abbreviations

ECM: extracellular matrix
ENS: enteric nervous system
ICC: interstitial cell of Cajal
PαC: PDGFRα+ cell
SIP syncytium: <u>S</u>mooth muscle Interstitial cells of Cajal and <u>P</u>DGFRα+ cell Syncytium
SMC: smooth muscle cell

## Supplemental Figures and Tables

**Supplementary Table S1:** Characteristics of human colon samples. Additional details are provided in Tables 2 and Table 3 in Wright and Schneider *et al.* 2021 ^18^.

| Sample ID (GEO accession number) | Age | Sex | Surgical Indication | Colon region | Number of barcoded cells per sample |
| --- | --- | --- | --- | --- | --- |
| 4579 (GSM4747491) | 54 | M | Cecal polyp | Right colon | 680 |
| 4602 (GSM4747492) | 75 | M | Hx of cecal lesion | Right colon | 2,316 |
| 4683 (GSM4747493) | 38 | M | Goblet cell carcinoma | Right colon | 2,081 |
| 4695 (GSM4747494) | 77 | F | Colonic mass | Right colon | 833 |
| 4701 (GSM4747495) | 78 | M | Rectal cancer | Sigmoid colon | 4,414 |
| 4950 (GSM4747496) | 78 | M | Bowel obstruction | Sigmoid colon | 1,237 |
| 4969 (GSM4747497) | 83 | M | Adenocarcinoma | Right colon | 653 |
| 4966 (GSM4747498) | 71 | F | Bowel obstruction | Right colon | 432 |
| 4988 (GSM4747499) | 65 | F | Colon polyp | Right colon | 237 |
| 4992 (GSM4747500) | 47 | M | Rectal carcinoma | Sigmoid colon | 60 |
| 5031 (GSM4747501) | 70 | M | Colon polyp | Right colon | 1728 |
| 5035 (GSM4747502) | 24 | M | Volvulus | Sigmoid colon | 2524 |
| 5040 (GSM4747503) | 44 | M | Colonic mass | Right colon | 755 |
| 5047 (GSM4747504) | 65 | M | Rectal<br>adenocarcinoma | Sigmoid<br>colon | 957 |
| 5054 (GSM4747505) | 36 | F | Bowel adhesions | Right colon | 1219 |
| 5059 (GSM4747506) | 59 | F | Adenocarcinoma | Right colon | 338 |

**Supplementary Table S2:** Identification of clusters based on gene expression.

| Cell Type | Defining genes |
| --- | --- |
| B Cells | *BLK, IGHM, IGHD, CD79A* |
| Fibroblasts | *MEOX2, VIM, COL1A1, COL1A2, FMO1, LSP1<sup>58</sup>* |
| Glia | *S100B, SOX10* |
| Glia#1 | *PLP1, PMP22, S100B* |
| Glia#2 | *VIP, GAL, S100B, PLP1, DSCAM, ELAVL4* |
| Interstitial Cells of Cajal (ICCs) | *ANO1, KIT* |
| Lymphatic Endothelial Cells | *FLT4, LYVE1, CD36, PECAM1, CLDN5, CCL21, TIE1* |
| Macrophages | *CD14, CD68, CXCR4, CD163, MSR1, LYZ* |
| Mast Cells | *KIT, CD36, HDC* |
| Myofibroblasts | *FAP, DES, AOC3, TPM2, TNC, MYOCD, MKL1, RYR2* |
| PDGFRα#1 (PaC#1) | *PDGFRA, KCNN3, P2YR1* |
| PDGFRα#2 (PaC#2) | *PDGFRA, KCNN3, LAMA2, FBLN1, FBN1, TNXB, COL1A1, COL1A2, COL6A3, MMP2, TIMP1* |
| Smooth Muscle Cells (SMC) | *ACTG2, ACTA2, MYH11, MYLK, MYOCD, TAGLN* |
| T Cells | *CD2, CD3D, CD52, GZMA, TRAC* |
| Vascular Endothelial Cells | *PECAM1, FLT1, VWF, FABP4, CDH5, TIE1* |

